# Large adipocytes alter mode of lipid release and promote breast cancer malignancy

**DOI:** 10.1101/2025.03.28.645549

**Authors:** Garrett F Beeghly, Marlee I Pincus, Rohan R Varshney, Dilip D Giri, Domenick J Falcone, Michael C Rudolph, Marc A Antonyak, Neil M Iyengar, Claudia Fischbach

**Affiliations:** Nancy E. and Peter C. Meinig School of Biomedical Engineering, Cornell University, Ithaca, NY, USA; Harold Hamm Diabetes Center and Department of Physiology, University of Oklahoma Health Sciences Center, Oklahoma City, OK, USA; Department of Pathology, Memorial Sloan Kettering Cancer Center, New York, NY, USA; Department of Pathology and Laboratory Medicine, Weill Cornell Medical College, New York, NY, USA; Department of Molecular Medicine, Cornell University, Ithaca, NY, USA; Department of Medicine, Memorial Sloan Kettering Cancer Center, New York, NY, USA; Department of Medicine, Weill Cornell Medical College, New York, NY, USA; Kavli Institute at Cornell for Nanoscale Science, Cornell University, Ithaca, NY, USA

## Abstract

Primary adipocytes possess a dramatic capacity to expand and retract in volume, leading to high variability in cell size within and between individuals. Yet, how adipocyte size impacts cell function remains unclear as adipocyte size is not tunable with traditional experimental approaches, forcing previous work to rely on correlative studies. Here, we develop protocols to separate primary adipocytes from the same donor into large and small populations and maintain these size-sorted cells in culture. Using these methods, we perform transcriptomic, lipidomic, and functional analyses on large and small adipocytes across two orthogonal mouse models of obesity and validate our results with human clinical samples. Our findings indicate that changes to cell size, rather than global differences mediated by weight gain, drive the transcriptional response of primary adipocytes to obesity. Moreover, large adipocytes shift from a traditional, lipase-mediated mode of lipid release to a non-canonical, extracellular vesicle-mediated mechanism. In functional coculture studies, this change promotes lipid accumulation in neighboring breast cancer cells, increasing their migration and proliferation via enhanced tumor cell fatty acid oxidation. Consistent with our experimental data, human patients with large adipocytes present with greater rates of dyslipidemia and higher concentrations of fasting triglycerides, even when accounting for differences in body mass index. Collectively, our results provide direct evidence that large and small adipocytes from the same donor differ in gene expression, lipid composition, and function with implications for the management of adipose tissue-related pathologies such as breast cancer.

## Introduction

White adipose tissue (WAT) expands via a combination of hyperplasia (increase in adipocyte number) and hypertrophy (increase in adipocyte volume)^1^, leading to high variability in cell size within and between individuals. Increased adipocyte size is characteristic of obesity and correlates with WAT dysfunction and insulin resistance^2^. However, large adipocytes also exist in lean individuals and are especially prevalent in those with metabolic disorders^3,4^. Moreover, changes in adipocyte size also affect the prognosis of patients with other diseases. For example, increased adipocyte size correlates with the transition from non-invasive to invasive breast cancer regardless of a given patient’s body mass index (BMI)^5^. Yet, how adipocyte size impacts cell function remains largely unclear due to insufficient strategies to manipulate adipocyte subpopulations of interest in a tractable manner.

Indeed, most studies of adipocyte function extrapolate differences by comparing lean and obese tissue from animal models or humans, which contain cells of varied size. This approach also introduces additional variables (e.g., diet composition, genetic differences), making it difficult to attribute observations to adipocyte size alone. *In vitro* studies could address these limitations but typically rely on progenitor cells differentiated on two-dimensional substrates, which are smaller in size and functionally distinct compared to primary cells^6^. Primary adipocytes are particularly challenging to culture due to their distinct properties. Notably, primary adipocytes are large (often over 100 μm in diameter), buoyant in aqueous media, post-mitotic, and fragile, which renders them incompatible with many cell culture platforms^7^ and single-cell technologies^8^. Thus, improving our understanding of how adipocyte size impacts diseased states will require new experimental approaches that enable comparisons of differently sized adipocytes isolated from the same donor tissue.

Here, we develop protocols to decouple adipocyte size from other variables and overcome the limitations of traditional animal models and two-dimensional culture systems. We first establish our ability to sort primary adipocytes from the same donor tissue into large and small cell populations across both diet-induced and genetic mouse models of obesity. Using RNA sequencing and targeted lipidomics, we characterize inherent phenotypic differences between large and small adipocytes while controlling for other obesity-associated effects. Subsequently, we maintain these adipocytes in coculture^9^ with cancer cells *ex vivo* given the clinical connection between adipocyte size and poor prognosis for breast cancer patients^5^. We monitor tumor cell lipid accumulation, migration, proliferation, and oxidative metabolism as a function of coculture with different adipocyte subpopulations and assess potential means of intercellular communication which mediated our observations. We then validate the relevance of our findings with a cohort of human mastectomy patients^10^ for which banked histology, paired RNA sequencing, and pre-operative blood panels are available. Collectively, our results provide direct evidence that large and small adipocytes from the same donor differ in gene expression, lipid composition, and function with implications for the management of breast cancer and metabolic diseases more broadly.

## Results

### Hypertrophic adipocytes can be isolated by differential buoyancy and size exclusion

Primary adipocytes are specialized, terminally differentiated cells which contain mostly lipid by volume and vary substantially in size (**Fig. 1A**). Previous *in vitro* studies have failed to resolve size-dependent differences in adipocyte function due to a reliance on differentiated progenitor cells (**Fig. S1**), which are multilocular, smaller in size, and functionally distinct compared to primary adipocytes^6^. To address these challenges, we developed experimental strategies to isolate the effect of cell size on adipocyte function while accounting for other covariates associated with weight gain. First, we assessed the size distribution of adipocytes in both diet-induced and genetic mouse models of obesity and their respective lean controls. For diet-induced studies, female C57BL/6 mice were fed a high-fat diet (HFD) or low-fat diet (LFD) for 12 weeks starting at 8 weeks of age (**Fig. 1B**). For genetic studies, we used female B6.Cg-Lep^Ob^ mice (*ob/ob*) which become obese due to a loss-of-function mutation in the satiety hormone leptin compared to wild-type (WT) mice^11^ (**Fig. 1C**). Consistent with the development of obesity, HFD and *ob/ob* mice weighed significantly more compared to LFD and WT mice at the time of adipocyte isolation (**Fig. S2**). As expected, adipocytes in the fat pads of both HFD and *ob/ob* mice were larger relative to adipocytes in their respective lean controls (**Fig. 1D-E**). Of note, we found that adipocyte hypertrophy was more pronounced in *ob/ob* mice than HFD mice, likely due to the impaired capacity of *ob/ob* progenitor cells to undergo adipogenesis and thus hyperplastic expansion^12^. However, both HFD and *ob/ob* mice contained a broad distribution of cell sizes, including adipocytes similar in size to those found in LFD and WT mice in addition to hypertrophic cells characteristic of obesity. We recognized this inherent heterogeneity as an opportunity to examine the impact of cell size on adipocyte function while controlling for other systemic effects of obesity.

**Figure 1.**
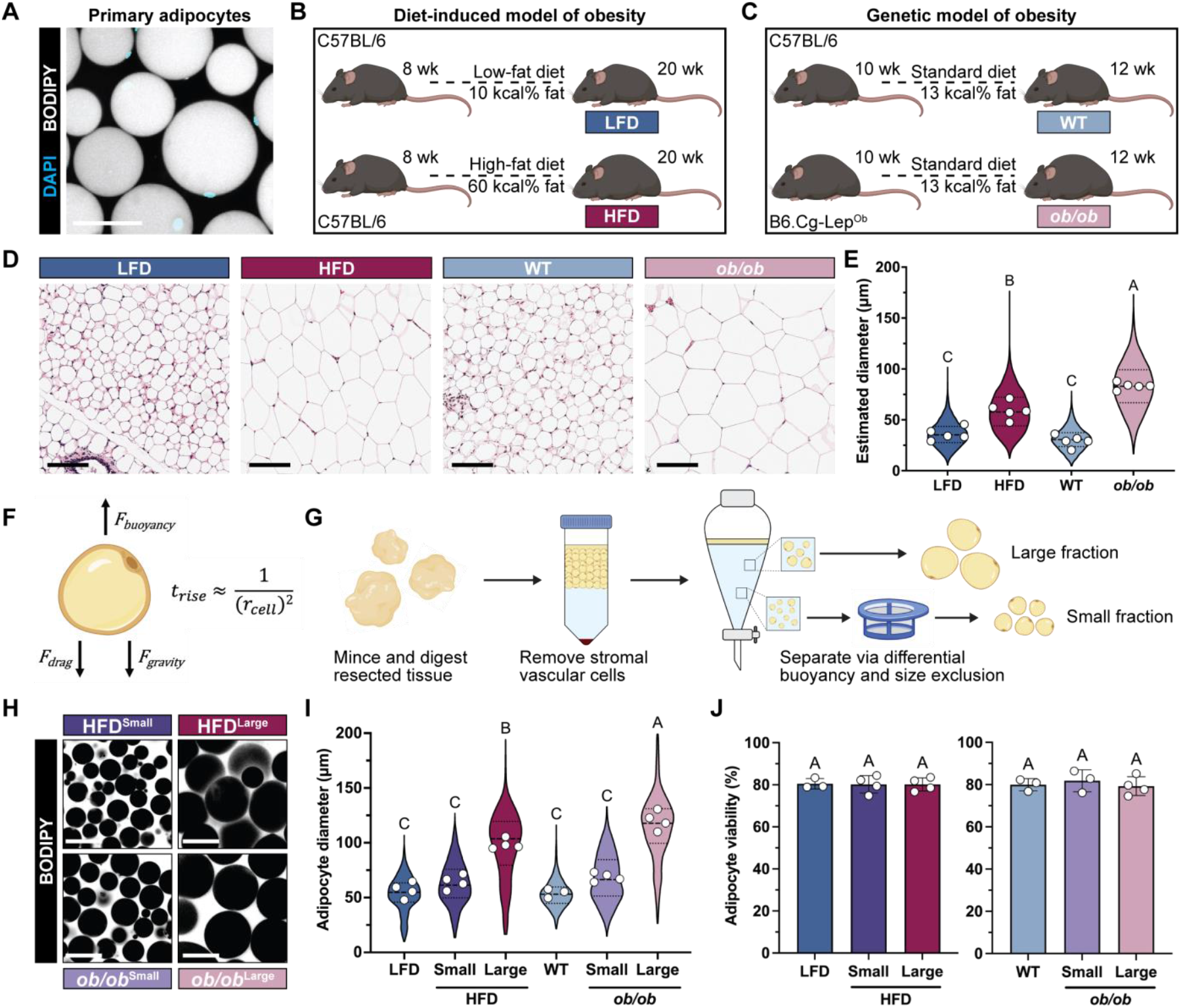
Hypertrophic adipocytes can be isolated by differential buoyancy and size exclusion. (**A**) Representative image of primary adipocytes stained for nuclei (cyan) and neutral lipid (gray). Scale bar = 100 μm. Schematic of (**B**) diet-induced and (**C**) genetic mouse models of obesity used to isolate primary adipocytes. (**D**) Hematoxylin- and eosin-stained sections of the anterior and inguinal fat pads of lean and obese mice. Scale bars = 100 μm. (**E**) Quantification of adipocyte diameters estimated from histology. Violin plots show the distribution of all quantified diameters and circles show the average adipocyte diameter per mouse (n = 5). (**F**) Free body diagram of the forces acting on a primary adipocyte in an aqueous solution. (**G**) Schematic of protocol to separate adipocytes by size. (**H**) Representative images of size-sorted adipocytes stained for neutral lipid (gray). Scale bars = 100 μm. (**I**) Quantification of sorted adipocyte diameters. Violin plots show the distribution of all quantified diameters and circles show the average adipocyte diameter per mouse (n = 3–4). Dashed line indicates the average lipid droplet diameter of *in vitro* differentiated adipocytes after two weeks for comparison. (**J**) Cell viability of lean and sorted obese adipocytes determined via propidium iodide exclusion (n = 3–4). All data are presented as mean ± standard deviation unless otherwise noted. Statistics were performed with a nested one-way analysis of variance (ANOVA) with Tukey’s multiple comparisons test. Compact letter display indicates statistical significance (p = 0.05).

To assess the impact of hypertrophy on adipocyte function independently of other obesity-associated changes, we separated adipocytes isolated from HFD and *ob/ob* mice into large and small cells by leveraging size-dependent differences in buoyancy^13^. Due to their lipid content, primary adipocytes experience a buoyancy force and rise out of aqueous solutions over time. This movement is opposed by a drag force due to fluid resistance. However, the buoyancy force scales with adipocyte volume while the drag force, via Stokes’ Law, scales with adipocyte diameter. Thus, the time for a given adipocyte to rise out of solution is inversely proportional to its radius squared (**Fig. 1F**). We exploited this phenomenon to separate heterogenous mixtures of adipocytes into cell populations of defined sizes (**Fig. 1G**). Notably, this approach enabled us to obtain small adipocytes from HFD and *ob/ob* mice that were comparable in diameter to adipocytes isolated from LFD and WT mice, respectively (**Fig. 1H-I**). Consistent with histology, large cells from *ob/ob* mice were greater in size than large cells from HFD mice. We also confirmed that size sorting adipocytes from HFD and *ob/ob* mice did not negatively impact cell viability compared to unsorted adipocytes from LFD and WT mice (**Fig. 1J**). Indeed, we maintained viability around 80% for all primary adipocyte subpopulations, consistent with previous results from our lab and others^6,9^. Collectively, these findings confirm that we can reliably isolate small and large adipocytes across two orthogonal mouse models of obesity, enabling us to analyze the effect of cell size on adipocyte function independently of other factors.

### Large and small adipocytes are transcriptionally distinct

Due to their buoyancy and size, primary adipocytes are not compatible with traditional single-cell RNA sequencing technologies^7^. While single-nucleus RNA sequencing has demonstrated that distinct subpopulations of primary adipocytes exist in mice and humans^8^, this approach does not capture the majority of coding RNA (located in the cytoplasm) or resolve physical variables such as cell size. Thus, to broadly assess how size affects adipocyte phenotype, we first performed bulk RNA sequencing on size-sorted adipocytes from HFD and *ob/ob* mice and unsorted adipocytes from LFD and WT mice. Principal component analysis (**Fig. 2A**) and hierarchical clustering (**Fig. 2B**) grouped samples based on condition. For both models of obesity, small adipocytes clustered in between large adipocytes and adipocytes from their respective lean controls, which indicates adipocyte dysfunction in obesity may exist along a continuum. Interestingly, small HFD adipocytes clustered more closely to LFD and WT adipocytes than large HFD adipocytes, suggesting that changes to cell size rather than global differences due to diet drive the transcriptional response of adipocytes to obesity. Of note, adipocytes isolated from *ob/ob* mice clustered further away from adipocytes isolated in all other conditions likely due to transcriptional differences driven by global leptin deficiency. Indeed, this result is consistent with sequencing of unsorted adipocytes, which demonstrate a greater number of differentially expressed genes between *ob/ob* and WT adipocytes than HFD and LFD adipocytes (**Fig. S3**). When comparing large and small adipocytes from HFD and *ob/ob* mice, we found a hypertrophy-associated gene signature of 821 differentially expressed genes that were shared across both models of obesity (**Fig. 2C**). These results suggest that altered adipocyte size causes distinct transcriptional changes that are independent of how obesity develops (e.g., differences in diet composition for LFD versus HFD mice or genetic differences and quantity of diet consumed for WT versus *ob/ob* mice).

**Figure 2.**
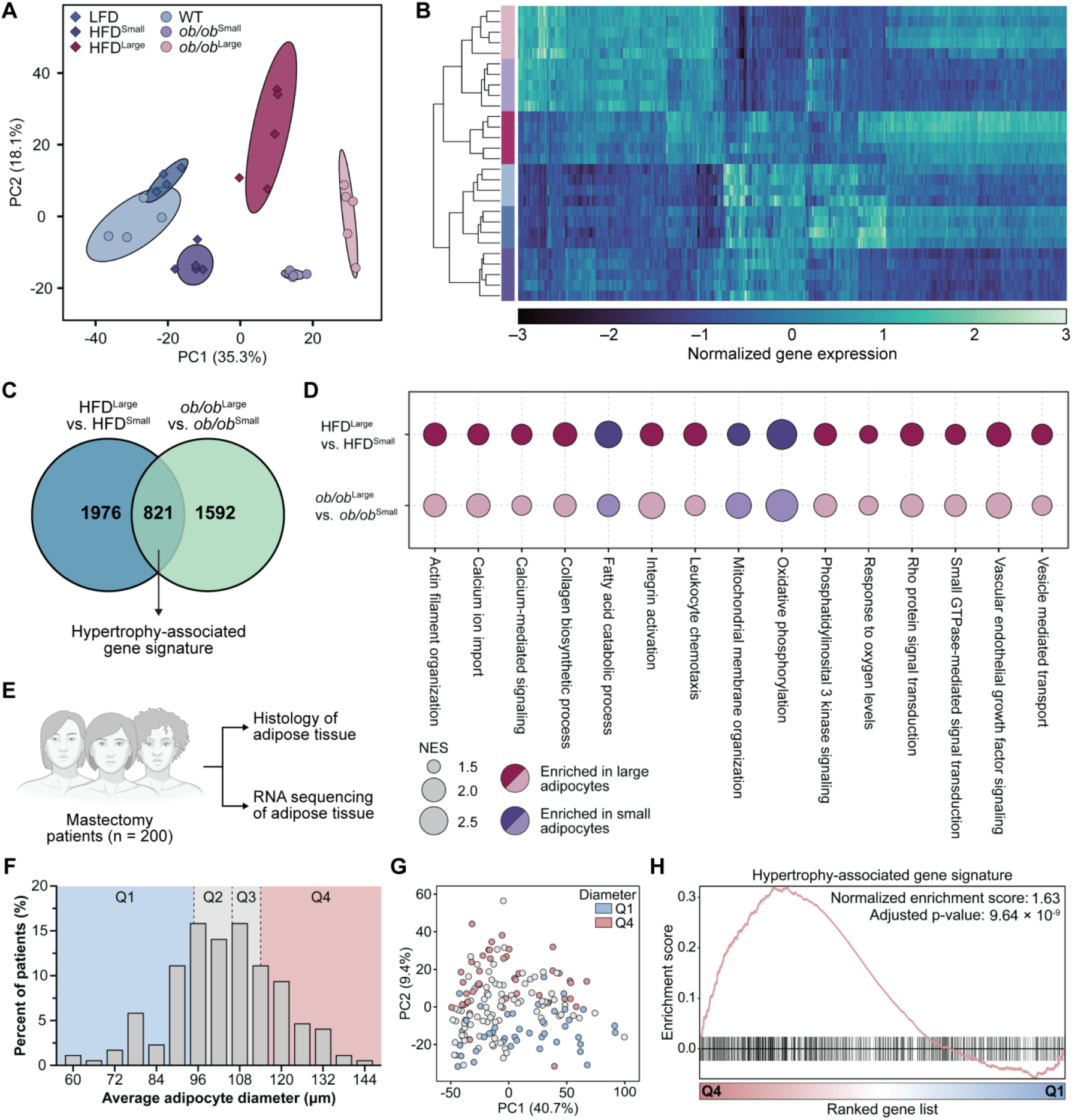
Large and small adipocytes are transcriptionally distinct. (**A**) Principal component analysis of gene expression data for lean adipocytes and size-sorted obese adipocytes. Ellipses show the 95% confidence interval for each condition and individual points represent replicates (n = 4–5). (**B**) Heatmap with hierarchical clustering of normalized gene expression data for the top 500 most variable genes between samples. (**C**) Venn diagram of differentially expressed genes (FDR < 0.05) between large and small adipocytes from HFD and *ob/ob* mice, including a hypertrophy-associated gene signature of 821 genes shared across both models. (**D**) Gene sets from Gene Ontology Biological Processes (GOBP) enriched between large and small adipocytes across both models. All comparisons shown meet the criteria set for statistical significance (FDR < 0.05). (**E**) Schematic of paired histology and RNA sequencing data from a cohort of 200 human mastectomy patients. (**F**) Histogram of patients stratified into quartiles based on their average adipocyte diameter determined from histology. (**G**) Principal component analysis of gene expression data for each patient. Circles are colored based on quartile classification. (**H**) Enrichment plot of patients in Q4 versus Q1 for the hypertrophy-associated gene signature defined in (C).

Next, we assessed the functional states associated with adipocyte hypertrophy for both obesity models using gene set enrichment analysis (GSEA). Large adipocytes from HFD and *ob/ob* mice were enriched for gene sets related to adipose tissue dysfunction in obesity^1^ including altered collagen deposition, lipid processing, pro-inflammatory signaling, response to hypoxia, and angiogenic signaling (**Fig. 2D**). Interestingly, we also found that large adipocytes were enriched for gene sets related to cytoskeleton organization, calcium- and integrin-mediated mechanosignaling, and RhoA signal transduction, consistent with recent studies indicating that adipocytes mechanically respond to cell expansion^14–17^. In contrast, small adipocytes were enriched for gene sets related to lipid catabolism, mitochondrial organization, and oxidative phosphorylation. These results are consistent with prior studies demonstrating that enlarged adipocytes exhibit defective aerobic metabolism due to mitochondrial fragmentation and dysfunction^18,19^. To assess the relevance of our findings in humans, we used a cohort of 200 mastectomy patients for which banked histology and paired RNA sequencing of uninvolved (tumor free) WAT were available^10^ (**Fig. 2E**). We first quantified the mean adipocyte diameter using histology for each patient to compare the transcriptomes of WAT with larger and smaller adipocytes on average (e.g., patients with adipocyte sizes in the fourth and first quartiles, respectively) (**Fig. 2F**). Patients in the fourth quartile of adipocyte diameter demonstrated global differences in gene expression (**Fig. 2G**) and were significantly enriched for the hypertrophy-associated gene signature we defined from murine adipocytes (**Fig. 2H**). Interestingly, patients in the fourth quartile of adipocyte diameter were also enriched for similar biological processes from GSEA as large cells in mice (**Fig. S4**). Collectively, these results indicate that large and small adipocytes are transcriptionally distinct across two mouse models of obesity, that large adipocytes are enriched for gene sets related to WAT dysfunction, that small adipocytes are enriched for gene sets related to normal mitochondrial metabolism even in obese conditions, and that key transcriptional changes associated with altered adipocyte size are shared between mice and humans.

### Large adipocytes promote breast cancer cell lipid accumulation, migration, and proliferation

Motivated by the correlation between adipocyte size and poor prognosis for breast cancer patients^5^, we next aimed to assess how large versus small adipocytes functionally impact breast cancer cell behavior by encapsulating size-sorted adipocytes for *ex vivo* culture using protocols we recently established^9^. This experimental model mimics the collagen-rich extracellular matrix of WAT^20,21^, achieves a cell volume fraction similar to native tissue, and maintains the viability of primary adipocytes in three-dimensional culture for extended periods of time. Moreover, this setup facilitates coculture studies with breast cancer cells by allowing for the exchange of soluble factors between adipocytes and tumor cells while enabling separation of both cell types for downstream analysis (**Fig. 3A**). Initial studies with four different breast cancer cell lines and unsorted adipocytes suggested that, regardless of cell line, tumor cells accumulate more lipid when cultured with adipocytes from *ob/ob* relative to WT mice (**Fig. S5**). Of the cell lines tested, MDA-MB-231s acquired the greatest amount of lipid, possibly due to associations between altered lipid metabolism and aggressive mesenchymal cell states^22^. Given these findings, MDA-MB-231s were selected for future studies. To account for relative differences in adipocyte size, we kept the total volume rather than total number of adipocytes constant between conditions. After three days of coculture, MDA-MB-231 breast cancer cells were either fixed and stained for analysis via fluorescence microscopy or reseeded for functional assays (**Fig. 3B**). By staining for neutral lipid with BODIPY 493/503 (**Fig. 3C, K**), we found that breast cancer cells cocultured with large adipocytes from HFD or *ob/ob* mice accumulated significantly more lipid than cancer cells cocultured with small adipocytes from HFD or *ob/ob* mice, adipocytes from LFD or WT mice, or monocultured controls (**Fig. 3D, L**). Across both models of obesity, this increase in neutral lipid was due to a greater number of lipid droplets per cell as well as a greater average lipid droplet size (**Fig. 3E, M**), suggesting that large adipocytes promote lipid uptake or biosynthesis by neighboring cancer cells.

**Figure 3.**
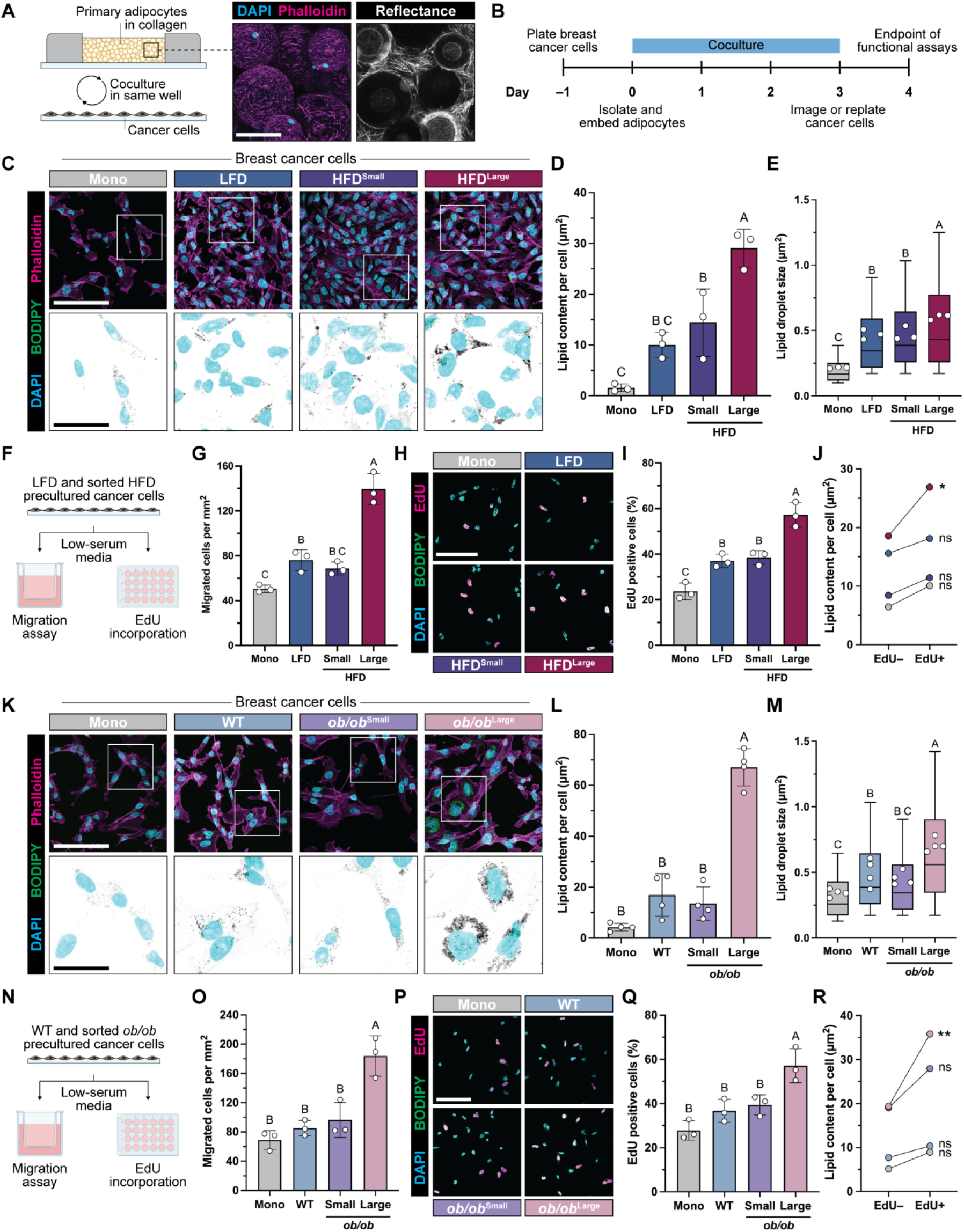
Large adipocytes promote lipid accumulation, migration, and proliferation of breast cancer cells. (**A**) Schematic of coculture setup between primary adipocytes and MDA-MB-231 breast cancer cells with a representative image of collagen-embedded adipocytes stained for nuclei (cyan) and F-actin (magenta). Confocal reflectance image (right) shows collagen fiber structure between cells. Scale bar = 100 μm. (**B**) Timeline of coculture experiments. Representative images of cancer cells after monoculture (mono) or coculture with lean or size-sorted obese adipocytes for diet-induced (**C**) and genetic (**K**) mouse models of obesity. Scale bars = 100 μm. Insets show nuclei (cyan) and neutral lipid (gray). Inset scale bars = 35 μm. Quantification of neutral lipid staining with BODIPY 493/503 per cancer cell for diet-induced (**D**) and genetic (**L**) models (n = 3–4). Size distribution of quantified lipid droplets for diet-induced (**E**) and genetic (**M**) models. Box and whisker plots show the distribution of all droplets and circles show the average droplet size per replicate (n = 3–4). Schematic of migration and proliferation assays of precultured cancer cells for diet-induced (**F**) and genetic (**N**) models. Number of migrated cancer cells 12 hours after reseeding onto tissue-culture inserts for diet-induced (**G**) and genetic (**O**) models (n = 3). Representative images of cancer cells 14 hours after reseeding and 2 hours after EdU treatment for diet-induced (**H**) and genetic (**P**) models. Quantification of EdU-positive cancer cells as a percent of total cells for diet-induced (**I**) and genetic (**Q**) models (n = 3). Average lipid content for EdU– and EdU+ cancer cells across diet-induced (**J**) and genetic (**R**) models (n = 3). All data are presented as mean ± standard deviation unless otherwise noted. Statistics for (J) and (R) performed with a paired t-test per condition. All other statistics performed with a one-way analysis of variance (ANOVA) with Tukey’s multiple comparisons test. Compact letter display indicates statistical significance (p = 0.05).

For functional studies, breast cancer cells precultured with size-sorted adipocytes were reseeded into collagen-coated tissue culture inserts for migration assays or onto glass coverslips for proliferation assays (**Fig. 3F, N**). We found that cancer cells precultured with large adipocytes from HFD or *ob/ob* mice migrated significantly more than cancer cells precultured with small adipocytes from HFD or *ob/ob* mice, adipocytes from LFD or WT mice, or monocultured controls. (**Fig. 3G, O**). Likewise, breast cancer cells precultured with large adipocytes from HFD or *ob/ob* mice were more proliferative relative to all other conditions as measured by incorporation of the thymidine analog 5-ethynyl-2’-deoxyuridine (EdU) (**Fig. 3H-I, P-Q**). Interestingly, EdU-positive cancer cells had more lipid than EdU-negative cancer cells on average across all conditions, but this difference was most pronounced and only statistically significant for breast cancer cells precultured with large adipocytes from HFD or *ob/ob* mice (**Fig. 3J, R**). These results suggest that increased cancer cell proliferation due to preculture with large adipocytes may be mediated by the greater presence of lipid in these cells, and we aimed to test this hypothesis moving forward. Collectively, our findings indicate that soluble factors released by large versus small adipocytes promote breast cancer cell lipid accumulation, as well as migration and proliferation. Since we observed similar trends between experiments performed with HFD and *ob/ob* mice, we focused our remaining studies on adipocytes isolated from *ob/ob* mice given that these mice do not require expensive and lengthy feeding periods prior to tissue isolation.

### Large adipocytes alter breast cancer cell lipid metabolism

Given that adipocytes are implicated in altering lipid metabolism in melanoma cells^23,24^ and that we observed adipocyte size positively correlated with lipid accumulation in cocultured breast cancer cells (**Fig. 3D, L**), we speculated that adipocyte size promoted tumor cell migration and proliferation in our studies by affecting lipid metabolism. Interestingly, fluorescence staining revealed that lipid droplets were closely associated with the mitochondrial network of MDA-MB-231 breast cancer cells cocultured with adipocytes (**Fig. 4A**), suggestive of altered lipid metabolism given that fatty acids (FAs) must be transported from cytoplasmic lipid droplets into the mitochondria to generate ATP via FA oxidation (FAO). To test this possibility, we first measured the oxygen consumption rate of breast cancer cells precultured with different adipocyte populations as a relative metric of mitochondrial activity in the presence or absence of etomoxir, which prevents FAO by irreversibly inhibiting carnitine palmitoyltransferase-1 (CPT1A) and precluding the transport of FAs into the mitochondria (**Fig. 4B**). At baseline, breast cancer cells precultured with large *ob/ob* adipocytes consumed more oxygen than those precultured with small *ob/ob* adipocytes, WT adipocytes, or monocultured controls as determined by metabolic flux analysis (**Fig. 4C**). Treatment with etomoxir reduced oxygen consumption rates for all precultured conditions to levels comparable to monocultured controls for both basal respiration (**Fig. 4D**) and maximal respiration (**Fig. 4E**), which was induced by the addition of the electron transport chain uncoupler FCCP. However, etomoxir had the greatest relative effect on MDA-MB-231 cells that were precultured with large *ob/ob* adipocytes. These results indicate that cell-cell communication between large adipocytes and breast cancer cells increases tumor cell oxygen consumption in a manner dependent on the oxidation of FAs as metabolic substrates. To determine if this increased mitochondrial activity was responsible for our observed changes in breast cancer cell behavior in our previous preculture studies, we repeated these experiments but treated cancer cells with or without etomoxir when replating for functional assays (**Fig. 4F**). Strikingly, the addition of etomoxir prevented increases to MDA-MB-231 migration (**Fig. 4G**) and proliferation (**Fig. 4H**) following preculture with large *ob/ob* adipocytes compared to vehicle controls. Etomoxir treatment also negated the correlation between lipid content and EdU incorporation (**Fig. 4I**), which suggests that an inability to use FAs as metabolic substrates prevents the growth advantage otherwise conferred to breast cancer cells with greater lipid accumulation. Collectively, our results indicate that large adipocytes secrete factors which increase the mitochondrial activity of breast cancer cells by promoting FAO and, in turn, facilitate their migration and proliferation.

**Figure 4.**
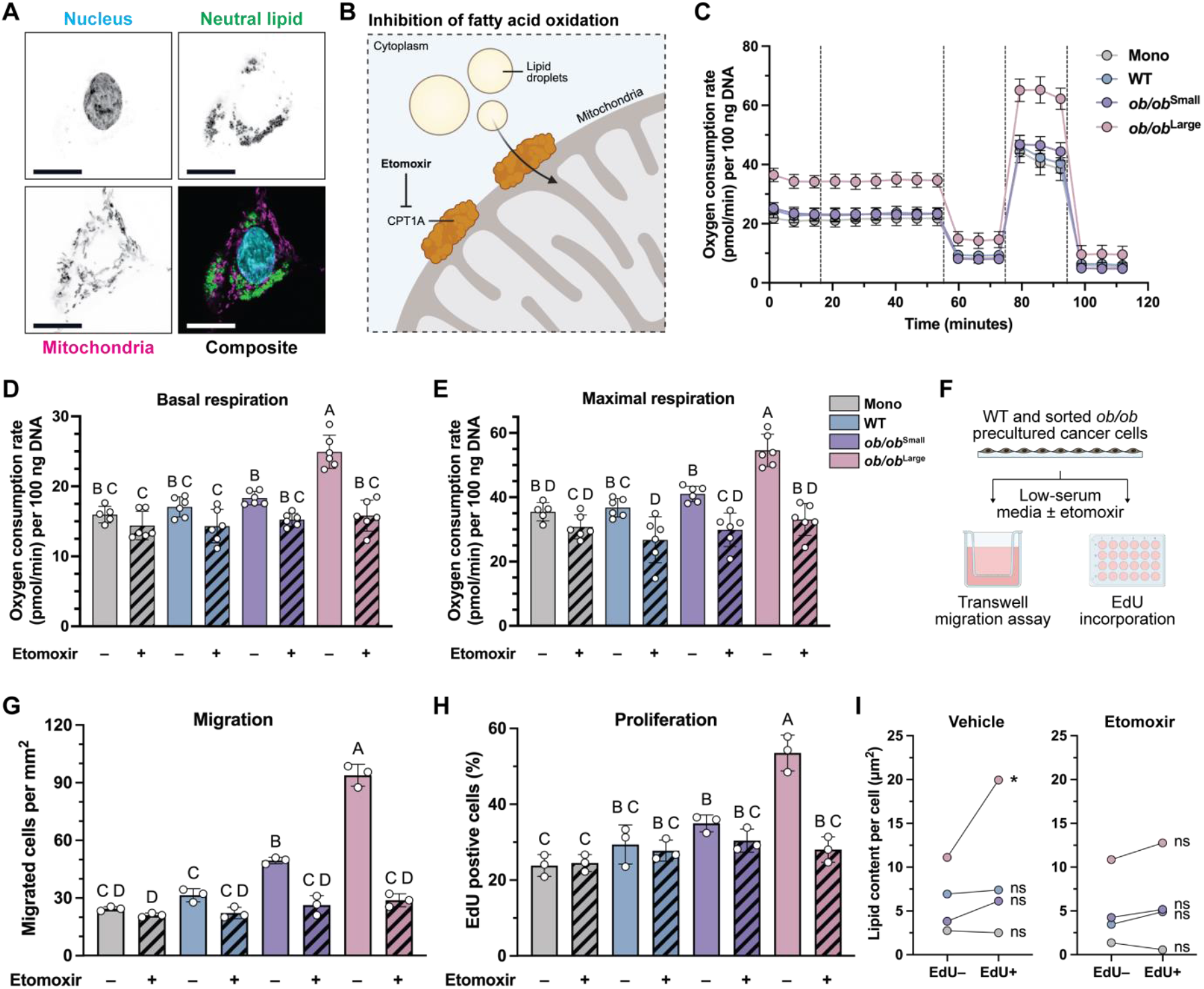
Large adipocytes alter breast cancer cell lipid metabolism to promote migration and proliferation. (**A**) Representative image of a precultured MDA-MB-231 breast cancer cell showing colocalization of neutral lipid droplets and mitochondria. Scale bar = 15 μm. (**B**) Schematic of FAO inhibition by etomoxir. (**C**) Normalized oxygen consumption rate of monocultured or precultured cancer cells over time. Dashed lines indicate sequential addition of etomoxir or vehicle, oligomycin, FCCP, and rotenone/antimycin A (n = 5–6). Only vehicle conditions are plotted to improve visibility. (**D**) Quantification of basal respiration rate per condition from (C) (n = 5–6). (**E**) Quantification of maximal respiration rate per condition from (C) (n = 5–6). (**F**) Schematic of migration and proliferation assays for precultured cancer cells treated with etomoxir or vehicle control. (**G**) Number of migrated cancer cells 12 hours after reseeding onto tissue-culture inserts with or without etomoxir (n = 3). (**H**) Quantification of EdU-positive cancer cells as a percent of total cells with or without etomoxir (n = 3). (**I**) Average lipid content for EdU– and EdU+ cancer cells across conditions (n = 3). All data are presented as mean ± standard deviation unless otherwise noted. Statistics for (I) performed with a paired t-test per condition. All other statistics performed with a one-way analysis of variance (ANOVA) with Tukey’s multiple comparisons test. Compact letter display indicates statistical significance (p = 0.05).

### Large adipocytes increase lipid transfer via extracellular vesicle-mediated release

We next aimed to assess if the increased lipid content of breast cancer cells cocultured with large adipocytes could be due to size-dependent differences in lipid release. To first confirm that cancer cells uptake lipid released by adipocytes, we prelabeled adipocytes with BODIPY 493/503 prior to coculture. After 48 hours, we observed the presence of prelabeled, adipocyte-derived lipid in cocultured breast cancer cells (**Fig. 5A**). These results suggest that adipocytes increase the lipid content of cocultured cancer cells via direct transfer of lipid rather than by changing tumor cell lipogenesis. We then aimed to determine how large adipocytes increase lipid transfer given that adipocytes can release lipids via two separate mechanisms. Canonical lipolysis by adipocytes is mediated by lipase-dependent breakdown of triglycerides into glycerol and non-esterified FAs, which then bind to carrier proteins for export into the extracellular space (**Fig. 5Bi**). In addition, adipocytes can release lipid via a lipase-independent mechanism where small lipid droplets containing intact triglycerides bud from the central lipid droplet and are packaged into extracellular vesicles for secretion^25^ (**Fig. 5Bii**). Conversely, cancer cells store triglycerides in their own lipid droplets, which requires esterification of FAs following uptake or lipogenesis (**Fig. 5C**). To determine which of these two mechanisms facilitates increased accumulation of lipid in breast cancer cells due to coculture with large adipocytes, we first measured the glycerol content of media collected from our experiments as a relative metric of lipolysis. Media collected from cocultures of cancer cells with large *ob/ob* adipocytes contained less glycerol than cocultures with small *ob/ob* or WT adipocytes (**Fig. 5D**). These results suggest that the rate of lipolysis, as estimated by glycerol release, cannot explain the observed increase in lipid content of cancer cells cocultured with large adipocytes. Therefore, we next tested whether adipocytes transfer lipid to cancer cells via extracellular vesicles and if this transfer differs as a function of adipocyte size. To this end, we purified extracellular vesicles from the media of a constant volume of small or large adipocytes in monoculture via ultracentrifugation for downstream analysis (**Fig. 5E**). We found that large *ob/ob* adipocytes secrete significantly more vesicles than small *ob/ob* or WT adipocytes (**Fig. 5F**). This increase was primarily driven by vesicles less than 200 nm in size, suggestive of greater exosome release (**Fig. 5G**). Moreover, we confirmed the purity of our isolated vesicles by blotting for the cell-specific marker FAK and the general extracellular vesicle marker FLOT2 (**Fig. 5H**). We then validated these results with adipocytes isolated from HFD and LFD mice. Similarly, we found that media from cocultures of cancer cells with large HFD adipocytes contained less glycerol than other conditions and that large HFD adipocytes secreted significantly more vesicles per cell volume than small HFD or LFD adipocytes (**Fig. S6**).

**Figure 5.**
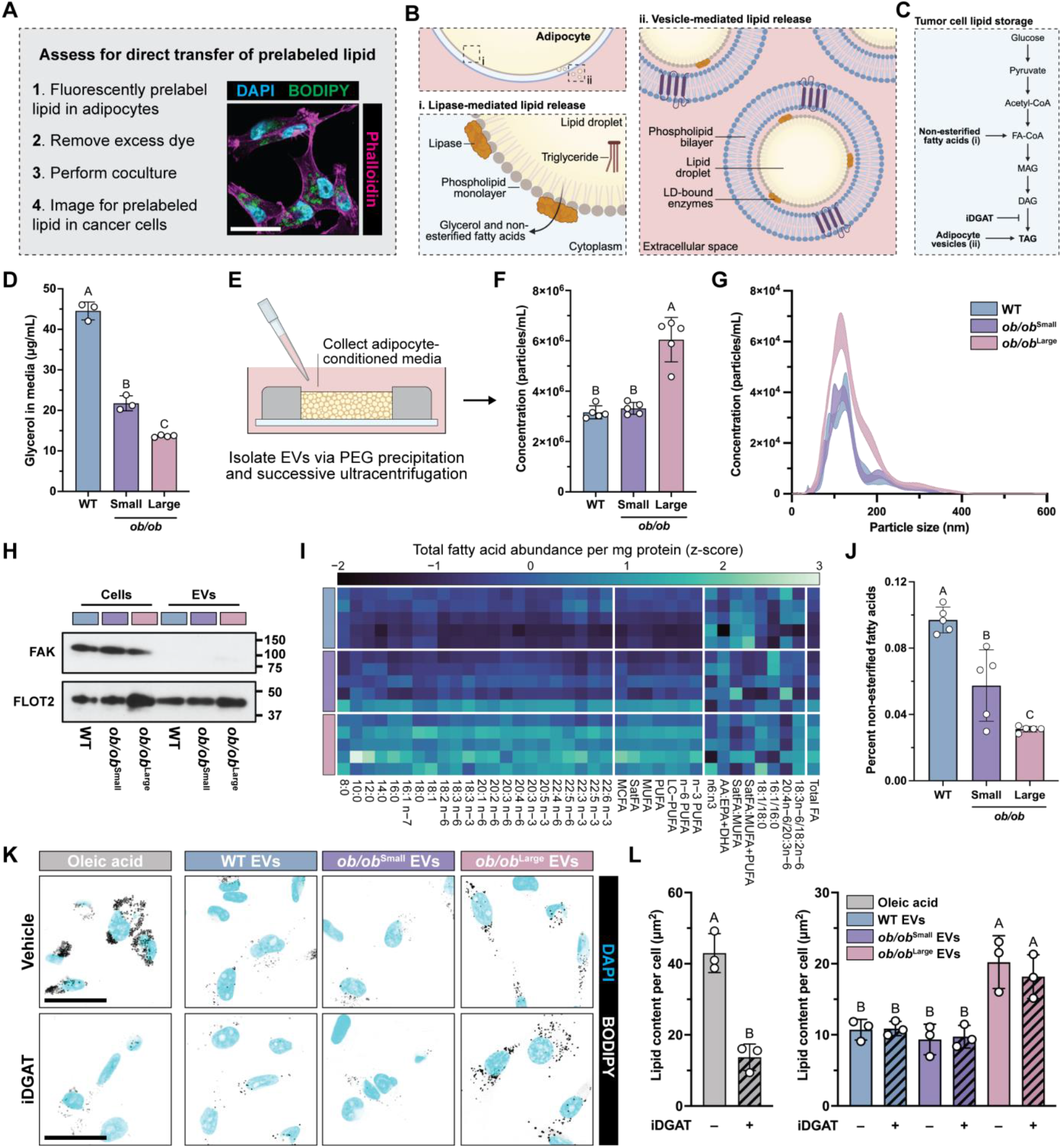
Large adipocytes promote breast cancer cell lipid accumulation through extracellular vesicle-mediated transfer of triglycerides. (**A**) Schematic of lipid transfer experiment. Adipocytes were pre-labeled with BODIPY 493/503 and washed to remove excess dye prior to the initiation of coculture. Scale bar = 35 μm. (**B**) Schematic of (i) lipase-mediated and (ii) vesicle-mediated lipid release by adipocytes. (**C**) Schematic of lipid storage by cancer cells. (**D**) Concentration of glycerol in conditioned media collected from cancer cells and adipocytes in coculture after 72 hours (n = 3–4). (**E**) Schematic of extracellular vesicle isolation from adipocyte-conditioned media. (**F**) Concentration of extracellular vesicles collected from adipocytes in monoculture after 24 hours (n = 5). (**G**) Size distribution of extracellular vesicles from (F) (n = 5). (**H**) Western blot of cell and extracellular vesicle lysates collected from lean and size-sorted obese adipocytes. (**I**) Heatmap of targeted lipidomics data showing the total FA composition of lean and size-sorted obese adipocytes normalized to protein content (n = 5). (**J**) Percent of non-esterified relative to total FAs from (I) (n = 5). (**K**) Representative images of MDA-MB-231 breast cancer cells 24 hours after treatment with oleic acid as a positive control or extracellular vesicles collected from WT, small *ob/ob*, or large *ob/ob* adipocytes. Cancer cells were also treated concurrently with vehicle or dual inhibitors for DGAT-1 and DGAT-2 (iDGAT). Scale bars = 35 μm. (**L**) Quantification of neutral lipid staining with BODIPY 493/503 from (K) per cancer cell (n = 4). All data are presented as mean ± standard deviation unless otherwise noted. Statistics were performed with a one-way analysis of variance (ANOVA) with Tukey’s multiple comparisons test. Compact letter display indicates statistical significance (p = 0.05).

To determine if adipocytes of varying size differed in their lipid composition which could, in turn, influence lipid accumulation in cocultured breast cancer cells, we performed targeted lipidomics of total and non-esterified FAs in WT, small *ob/ob*, and large *ob/ob* adipocytes (**Fig. 5I**). While large *ob/ob* adipocytes contained more saturated, unsaturated, and total FAs per mg protein relative to small *ob/ob* and WT adipocytes as expected, we did not observe consistent differences in specific lipid species in small versus large cells (**Fig. S7-S8**). However, large adipocytes contained a significantly lower percentage of non-esterified to total FAs compared to small adipocytes (**Fig. 5J**), consistent with our glycerol release data which suggested that large cells are less lipolytic relative to small cells. Interestingly, both small and large *ob/ob* adipocytes contained a higher ratio of arachidonic acid to eicosapentaenoic and docosahexaenoic acids than WT adipocytes, indicative of a pro-inflammatory lipid profile and suggestive of altered FA desaturase activity given that *ob/ob* and WT mice received the same diet (**Fig. S7-S8**). These results underscore the importance of examining adipocytes from the same donor when investigating the impact of size on adipocyte function, as some differences in adipocyte phenotype are donor-dependent regardless of size. Finally, to confirm that the increased lipid content of cancer cells cocultured with large versus small *ob/ob* adipocytes was due to vesicle-mediated transfer, we applied extracellular vesicles collected and purified from a constant volume of size-sorted adipocytes in monoculture to cancer cells in monoculture. Moreover, we inhibited DGAT-1 and DGAT-2 (iDGAT) concurrently with vesicle treatment in a subset of samples to prevent esterification of free FAs and further rule out the contributions of lipogenesis by cancer cells (**Fig. 5C**). After 24 hours, we found that breast cancer cells treated with vesicles from large *ob/ob* adipocytes contained significantly more lipid than those treated with vesicles from small *ob/ob* or WT adipocytes (**Fig. 5K-L**). However, lipid uptake was significantly lower across all conditions compared to coculture studies, likely due to the shorter period of vesicle collection (24 hours) than coculture (72 hours) and loss of vesicles during sample processing. Nevertheless, iDGAT prevented lipid storage in tumor cells treated with exogenous oleic acid as a positive control as expected, while iDGAT treatment did not affect lipid transfer via adipocyte-derived vesicles (**Fig. 5K-L**), indicating that these vesicles contain intact triglycerides rather than non-esterified FAs and that altered lipogenesis does not significantly contribute to cancer cell lipid content in our coculture system. Collectively, our results indicate that large adipocytes promote breast cancer cell lipid uptake through extracellular vesicle-mediated transfer of intact triglycerides.

### Adipocyte size is more predictive of systemic lipid dysfunction than body mass index

Given that we observed greater lipid release by large adipocytes in our coculture studies, we next aimed to determine if adipocyte size correlated with broader, systemic metabolic changes in our human clinical samples. To this end, we cross-referenced pre-operative blood panels for each of the 200 mastectomy patients in our cohort (**Fig. 6A**). Again, we stratified patients into quartiles based on their average adipocyte diameter determined via paired histology (**Fig. 6B**). Using this approach, we found that average adipocyte diameter (**Fig. 6C**) more accurately stratified patients diagnosed with dyslipidemia than BMI category (**Fig. 6D**). Likewise, average adipocyte diameter correlated more significantly with fasting triglycerides (**Fig. 6E**) than BMI (**Fig. 6F**), even when their interaction effects were considered using multiple linear regression (**Fig. 6G**). Thus, adipocyte diameter may be a more accurate predictor of lipid dysfunction, which is an established risk factor for breast cancer^26^, even for women with normal BMI. Collectively, these findings suggest that in addition to releasing more lipid to cells locally, as observed in our coculture studies, large adipocytes may also increase lipid release into circulation with consequences for whole-body metabolism.

**Figure 6.**
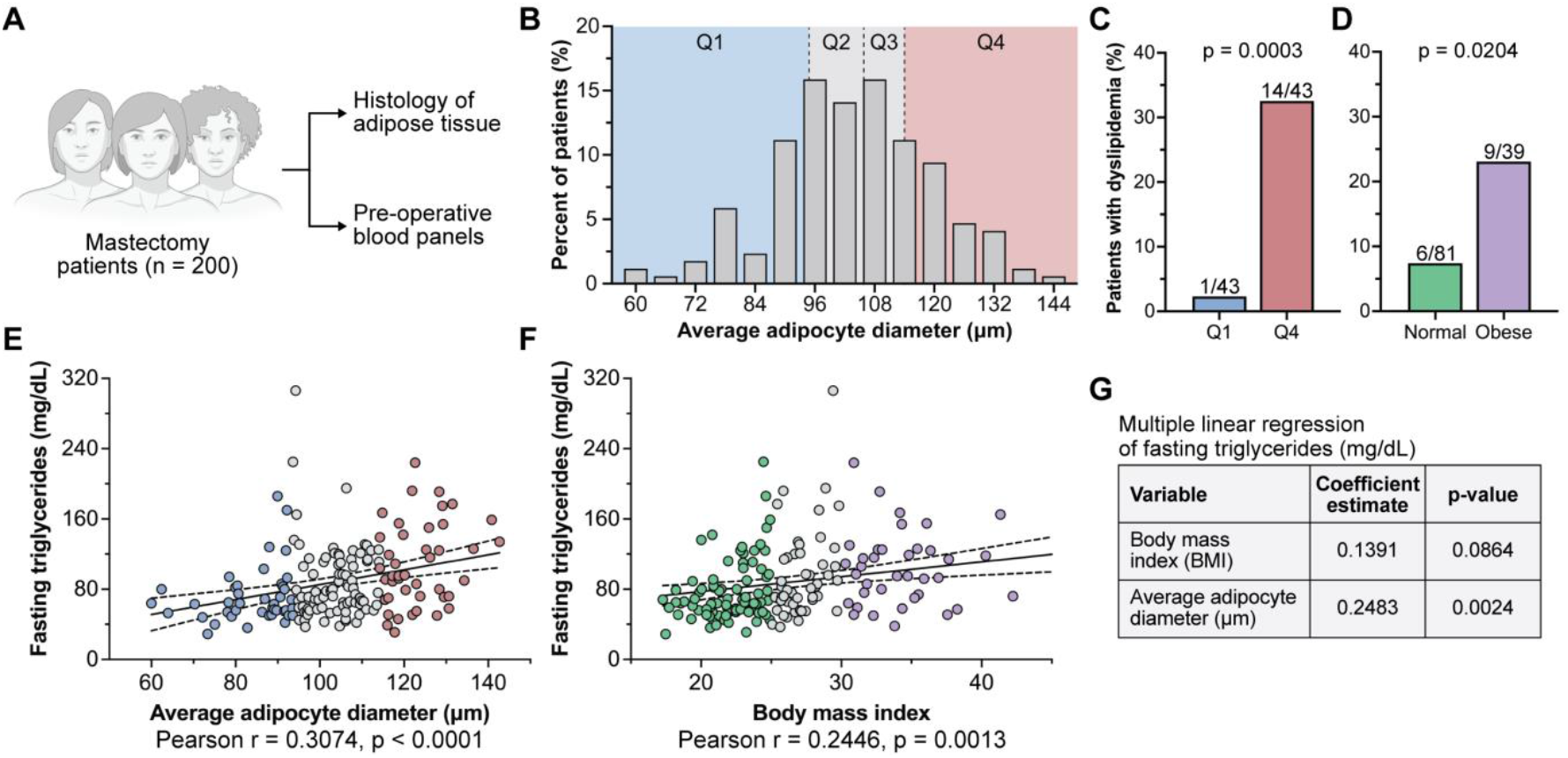
Adipocyte size is more predictive of systemic lipid dysfunction than body mass index. (**A**) Schematic of paired mammary adipose tissue histology and pre-operative blood panels from a cohort of 200 mastectomy patients. (**B**) Histogram of patients stratified into quartiles based on their average adipocyte diameter determined from histology. Percent of patients diagnosed with dyslipidemia when classified by (**C**) average adipocyte diameter or (**D**) BMI category (BMI < 25 for normal, BMI > 30 for obese). Statistics performed with a Fisher’s exact test per comparison. Simple linear regression of fasting triglycerides versus (**E**) average adipocyte diameter or (**F**) BMI. Statistics performed with a Pearson correlation analysis per comparison. (**G**) Multiple linear regression of fasting triglycerides as a function of both average adipocyte diameter and BMI following z-score data normalization.

## Discussion

How adipocyte size impacts cell function remains unclear due, in part, to limited methods to manipulate adipocyte subpopulations of interest. Our results, combining transcriptomic, lipidomic, and functional analyses, of size-sorted adipocytes provide direct evidence that large adipocytes are distinct from small adipocytes isolated from donor-matched tissue. Moreover, through extended culture of primary adipocytes *ex vivo*, we find that large adipocytes increase lipid accumulation in neighboring breast cancer cells through a non-canonical, extracellular vesicle-mediated mechanism of lipid release. In turn, large adipocytes enhance breast cancer cell migration, proliferation, and oxygen consumption in a manner dependent on FAO. Consistent with these experimental results, the presence of enlarged mammary adipocytes in human mastectomy patients correlated with systemic lipid dysfunction, even when controlling for differences in body mass index as a covariate. Collectively, our findings provide insight into how adipocytes respond to cell expansion not discernible with conventional approaches and further support the use of adipocyte size as a prognostic biomarker for breast cancer and metabolic disease.

Our understanding of adipocyte heterogeneity and biology remains limited given that single-cell analysis of primary cells is challenging due to their buoyancy and fragility^7,8^. While the use of single-nucleus RNA sequencing can overcome some of these limitations, this approach disregards cytoplasmic RNA, which constitutes the majority of coding RNA in cells, and fails to capture information about cell size. Here, we performed bulk RNA sequencing of primary adipocytes phenotypically sorted by size and identified that large, hypertrophic adipocytes constitute a transcriptionally distinct subpopulation of cells. Our findings are consistent with recent results demonstrating that WAT is more plastic^27,28^ and heterogenous^8,29^ than once considered. For example, multiple adipocyte subpopulations have been identified in mice with distinct sensitivities to insulin^29^ and whose relative proportions change with administration of a high-fat diet^8^. In our data, large adipocytes from both diet-induced and genetic mouse models of obesity were enriched for gene sets related to WAT dysfunction^1^, as were tissue samples collected from mastectomy patients with increased adipocyte size regardless of BMI. Moreover, large adipocytes also exhibited enrichment for gene sets related to collagen deposition, actin cytoskeleton remodeling, calcium ion transport, and Rho protein signal transduction, consistent with an emerging understanding of the mechanoresponsive adaptations required of adipocytes during drastic cell expansion^15,17^. To what extent these transcriptional changes are adipocyte intrinsic or a function of environmental cues in hypertrophic regions of WAT remains to be determined. Re-examining the transcriptome of large adipocytes after extended *ex vivo* culture could help make this distinction. Likewise, assessing hypertrophic adipocytes in the absence of obesity, such as those present in lean individuals with metabolic disease^3^, could help identify the relative contribution of cell intrinsic factors. Regardless of the cause, our findings underscore the need for a more nuanced view of WAT beyond BMI that accounts for other parameters such as adipocyte size. Indeed, in blood panels collected from our cohort of mastectomy patients, average adipocyte diameter correlated more significantly with dyslipidemia and fasting triglycerides than BMI, even when their interaction effects were considered—findings that are broadly consistent with the increased release of lipid by large adipocytes in our coculture experiments.

Previous studies have demonstrated that adipocytes can influence the behavior of neighboring tumor cells, though this connection is most established for melanoma compared to other carcinomas that develop in WAT-rich stroma such as breast cancer. For example, researchers found that melanoma cells overexpress fatty acid transport protein 1 (FATP1), which facilitates the uptake of adipocyte-derived non-esterified FAs and promotes tumor invasion^24^. In a separate study, adipocyte-derived exosomes, which were enriched for proteins implicated in FAO, mediated the metabolic reprogramming of melanoma cells^23^. In the context of breast cancer, several studies have linked adipocyte-derived lipids to altered tumor cell metabolism^30,31^. However, these studies used *in vitro* differentiated progenitor cells and thus did not assess obesity- or size-dependent effects. Indeed, while we saw trends for cocultures with small adipocytes, we only observed significant changes in breast cancer cell phenotypes (i.e. lipid content, migration, proliferation, oxygen consumption) for cocultures with large adipocytes. This may be due to our experimental setup, as we performed shorter cocultures than some of the cited studies^24,30^ or may reflect underlying differences in adipocyte-tumor cell communication when using primary adipocytes. Moreover, our results corroborate previous findings that obese WAT secretes more extracellular vesicles per mg tissue than lean WAT^25^ and that this increase is driven by adipocytes rather than cells of the stromal vascular fraction^23^. Our findings indicate that the greater rate of extracellular vesicle release by obese WAT likely stems from the increased presence of hypertrophic adipocytes in obesity, a conclusion not previously possible with gross analysis of tissue explants or unsorted cells. Moving forward, future studies will be needed to assess the impact of these adipocyte-derived vesicles on other breast cancer subtypes and stromal cells, such as tissue-resident macrophages and fibroblasts, within the tumor microenvironment.

Across experiments, our findings suggest that large adipocytes shift toward favoring an extracellular vesicle-mediated mode of lipid release, though the lack of endocrine signals which stimulate lipolysis in our system may be a confounding factor in these observations. Indeed, across both mouse and human sequencing data, hypertrophic adipocytes and hypertrophic WAT demonstrated positive gene set enrichment for vesicle-mediated transport and negative gene set enrichment for lipid catabolism. While initially counterintuitive given the greater degree of lipid transfer observed between large adipocytes and breast cancer cells in coculture, these findings are consistent with a lipase-independent, vesicle-mediated mode of lipid release by hypertrophic cells. Direct transfer of adipocyte-derived triglycerides via extracellular vesicles as opposed to non-esterified FAs via lipolysis could offer several potential advantages to recipient tumor cells. Beyond serving as metabolic substrates for FAO, a steady supply of vesicle-delivered triglycerides may also provide a readily available source of FAs for membrane biosynthesis, lipid signaling, and intracellular trafficking. In particular, tumor cells undergoing rapid division exhibit increased demand for phospholipids to sustain plasma membrane expansion, and specific lipid classes such as monounsaturated and polyunsaturated FAs are necessary for membrane fluidity and lipid raft formation^32^. Our lipidomics data indicate that large adipocytes contain an increased abundance of these classes of FAs relative to small adipocytes. This finding suggests that vesicle-mediated lipid transfer could facilitate tumor progression not only by fueling oxidative metabolism but also by supplying key structural lipids for tumor growth. Future untargeted lipidomics of the specific lipid species enriched in adipocyte-derived vesicles, as well as the lipids transferred to recipient tumor cells, will be critical to dissect this potential dual role of FAs in breast cancer malignancy.

In addition to triglycerides, previous studies have demonstrated that extracellular vesicles secreted by adipocytes are also enriched for lipid-droplet-associated proteins required for FAO^23,25^. Thus, by increasing vesicle release, hypertrophic adipocytes may provide tumor cells not only with FAs for mitochondrial oxidation but also the molecular machinery needed to metabolize them. Future proteomic studies of vesicles derived from large versus small adipocytes would reveal if altered protein cargo contributed to our observed increase in FAO for breast cancer cells precultured with large adipocytes. Moreover, excess quantities of non-esterified FAs, especially trans- or saturated FAs, can induce lipotoxicity by activating cell stress responses^33^. Indeed, inhibiting the ability of tumor cells to esterify free FAs, and thus induce lipotoxicity, has been explored as a therapeutic strategy for hepatocellular carcinoma^34^. In contrast, the direct transfer of intact triglycerides would bypass the need for FA esterification and limit the potential lipotoxic effects on cancer cells. Conversely, controlled oxidation of FAs by tumor cells has been shown to mediate chemoresistance to platinating agents^35^, which are increasingly prescribed for patients with advanced or triple-negative breast cancer^36^. Thus, moving forward, the effect of specific adipocyte populations of interest on breast cancer chemoresistance should be explored. More broadly, this shift in the mode of lipid release by hypertrophic adipocytes likely produces systemic metabolic alterations by altering the bioavailability and biodistribution of lipids in circulation, as supported by our patient data.

In conclusion, our results demonstrate that large and small adipocytes from the same donor differ in gene expression, lipid composition, and function across two orthogonal mouse models with key findings validated in human samples. Notably, large adipocytes shift to a non-canonical, extracellular vesicle-mediated mechanism of lipid release which reprograms the metabolism of nearby breast cancer cells, in turn promoting their migration and proliferation. Thus, these data provide an explanation for how large adipocytes directly contribute to breast cancer malignancy^5^ and to the poor clinical outcomes seen across obese patients in the clinic^37^. Our findings further posit that adipocyte size could serve as an independent prognostic biomarker for breast cancer patients regardless of BMI. Determining adipocyte size is clinically tractable given that peritumoral WAT is often included in routine biopsies used for diagnosis and automated approaches exist to characterize adipocyte morphology^38,39^. Collectively, our findings provide insight into the role of hypertrophic adipocytes as a distinct cell population within adipose tissue with implications for the management of breast cancer and metabolic diseases more broadly.

## Resource availability

All source data and code will be provided in the final publication of this manuscript.

## Acknowledgments

We thank all members of the Fischbach Lab for valuable discussion and input during the preparation of this manuscript; the Weill Institute of Cell and Molecular Biology and David McDermitt for access to an ultracentrifuge; Rebecca Williams for assistance with confocal microscopy; Tina Abratte for assistance with metabolic flux experiments; Malin Lönn for assistance with developing the adipocyte size-sorting protocol; the Cornell Center for Animal Resources and Education (CARE), Matthew Whitman, and Jessie Gorges for assistance with animal studies; the Cornell Animal Health Diagnostic Core for assistance with paraffin embedding and sectioning; and the Cornell Statistical Consulting Unit for assistance with statistical analysis. We acknowledge financial support from the NSF DGE1650441 (G.F.B.), NCI F31CA278410 (G.F.B.), NCI R01CA259195 (C.F., M.A.A.) and R01CA276392 (C.F.), the Center on the Physics of Cancer Metabolism NCI 1U54CA210184 (C.F.), and the Cornell Engineering Learning Initiatives program (M.I.P.). We also acknowledge support from the Cornell Biotechnology Resource Center Genomics Facility (RRID:SCR_021727) and the Cornell Biotechnology Resource Center Imaging Facility funded by NYSTEM C029155, NIH S10OD018516, and NIH S10RR025502.

## Author contributions

G.F.B. and C.F. conceived and designed experiments. G.F.B. performed all experiments with assistance from M.I.P. unless otherwise noted. R.R.V. and M.C.R. performed targeted lipidomics and associated analysis. M.A.A. performed Western blotting of cell and extracellular vesicle lysates. D.D.G., D.J.F., and N.M.I. obtained and analyzed the cohort of human mammary adipose tissue collected from mastectomy patients. G.F.B. prepared all figures and schematics. G.F.B. and C.F. wrote the manuscript with input from all authors.

## Supplemental data

Figures S1–S8.

## Methods

### Culture of breast cancer cell lines

MDA-MB-231 (RRID:CVCL_0062), MCF7 (RRID:CVCL_0031), PY8119 (RRID:CVCL_AQ09), and EO771 (RRID:CVCL_GR23) breast cancer cell lines were maintained in high-glucose Dulbecco’s Modified Eagle Medium (Gibco #12800) supplemented with 10% (v/v) fetal bovine serum (FBS, R&D Systems #S11150) and 1% (v/v) penicillin-streptomycin (ThermoFisher #15070063) unless otherwise noted. Primary adipocytes and cocultures were maintained in a 1:1 mixture of low-glucose Dulbecco’s Modified Eagle Medium (Gibco #316000) and Ham’s F12 Nutrient Mix (Gibco #21700) supplemented with 10% (v/v) FBS and 1% (v/v) penicillin-streptomycin unless otherwise noted. For functional studies of precultured tumor cells, a 1:1 mixture of low-glucose Dulbecco’s Modified Eagle Medium and Ham’s F12 Nutrient Mix supplemented with 1% (v/v) FBS and 1% (v/v) penicillin-streptomycin was used. For experiments involving extracellular vesicle collection or treatment, exosome-depleted FBS (ThermoFisher #A2720803) was used to limit the presence of exogenous vesicles. All breast cancer cell lines were acquired from the American Type Culture Collection (ATCC) and screened for mycoplasma contamination using a PCR-based testing kit (SouthernBiotech #13100-01) according to manufacturer protocols.

### Maintenance and differentiation of 3T3-L1 preadipocytes

Prior to the induction of adipogenesis, 3T3-L1 preadipocytes (RRID:CVCL_0123) were seeded at a density of 5,000 cells per cm^2^ onto 12 mm glass coverslips coated with 30 μg/mL collagen type I (VWR #47747-218) and cultured until 80% confluent. Adipogenesis was initiated with induction media consisting of high-glucose DMEM supplemented with 5% FBS, 1% penicillin-streptomycin, 1 mM dexamethasone (Sigma-Aldrich #D4902), 500 mM IBMX (Krackeler Scientific #45-I7018), 20 mM indomethacin (Sigma-Aldrich #405268), and 200 μM insulin (Krackeler Scientific #45-91077C). After 48 hours, cells were changed to maintenance media consisting of high-glucose DMEM supplemented with 5% FBS, 1% penicillin-streptomycin, and 200 μM insulin. Maintenance media was replaced every 48 hours until experimental endpoints were reached. 3T3-L1 preadipocytes were acquired from the American Type Culture Collection (ATCC) and screened for mycoplasma contamination using a PCR-based testing kit (SouthernBiotech #13100-01) according to manufacturer protocols.

### Animal models

For all animal studies, WAT was isolated from the anterior and inguinal fat pads around the mammary glands of nulliparous female C57BL6 mice (The Jackson Laboratories). For the genetic model of obesity, wildtype C57BL6 mice (strain 000664) and leptin-deficient B6.Cg-Lep^ob^ mice (strain 000632) were purchased at 10 weeks of age and fed a standard rodent diet (LabDiet #5053) *ad libitum* until sacrifice at 12 weeks of age. For the dietary model of obesity, wildtype C57BL6 mice were purchased at 6 weeks of age and fed a standard rodent diet for a two-week acclimation period. At 8 weeks of age, mice were randomized and placed on either a low-fat diet (Research Diets #D12450J, 10 kcal% fat) or high-fat diet (Research Diets #D12492, 60 kcal% fat) *ad libitum* until sacrifice at 20 weeks of age. Diets were administered weekly and remaining chow from the previous week was discarded. All mice were housed between 21°C and 23°C. All animal protocols were approved by the Institutional Animal Care and Use Committees at Weill Cornell Medicine and Cornell University.

### Histology of resected adipose tissue

Pieces of resected WAT were collected from euthanized C57BL6 mice and fixed in 4% (w/v) paraformaldehyde in 1x PBS for 24 hours at 4°C. Resected tissue was then stored in 70% (v/v) ethanol in water at 4°C until processing. Samples were sent to the Cornell College of Veterinary Medicine Animal Health Diagnostic Center for paraffin embedding, sectioning, and staining with hematoxylin and eosin. Stained sections were imaged and digitized on an Aperio ScanScope CS2 (Leica) with a 40x objective.

### Primary adipocyte isolation

Resected WAT was immediately placed in cold DMEM/F12 until processing. Tissue was then minced for several minutes until separated into approximately 1 mm^3^ pieces. Floating tissue pieces were decanted and reserved while any remaining pieces were discarded. The reserved tissue was then digested in 1.5 mg/mL collagenase type I (Worthington Biochemical #LS004197) in Hank’s Buffered Salt Solution (Gibco #14065056) supplemented with 1% (w/v) bovine serum albumin (ThermoFisher #BP1600-100) in a vertical tube rotator for 50 minutes at 37°C. The resulting cell suspension was passed through 200 μm cell strainers (pluriSelect #435020003) to remove undigested pieces of tissue. Adipocytes were allowed to float out of solution for 10 minutes before the infranatant was aspirated and Krebs-Ringer HEPES buffer (116 mM NaCl, 25 mM HEPES, 4 mM KCl, 2 mM D-glucose, 1.8 mM CaCl_2_, 1 mM MgCl_2_) supplemented with 1% (w/v) bovine serum albumin (KRHB) was added. This process was repeated three times to remove contaminating stromal vascular cells. To ensure consistent cell seeding, isolated adipocytes were packed via centrifugation for 3 minutes at 100 g and the resulting layers of aqueous buffer and free lipid were aspirated before use as described^9^. All subsequent handling of isolated adipocytes was performed with wide bore pipette tips.

### Size sorting of primary adipocytes

To separate adipocytes by size, isolated cells were placed in a 125 mL separatory funnel (ThermoFisher #4301-0125) along with 50 mL of KRHB. The funnel was gently inverted several times to resuspend the adipocytes. After 35 seconds, the bottom 35 mL of cell suspension was eluted and passed through a 50 μm cell strainer (pluriSelect #435005003). 35 mL of fresh KRHB was added to the funnel and this process was repeated. Adipocytes that passed through the cell strainers were pooled and used as the “small” fraction for all experiments. 35 mL of fresh KRHB was then added to the funnel before gently inverting several times. After 15 seconds, the bottom 35 mL of cell suspension was eluted and discarded. This process was repeated two additional times to remove cells of intermediate size. The remaining adipocytes were treated as the “large” fraction for all experiments.

### Assessment of primary adipocyte viability

To assess cell viability after isolation and sorting, unfixed adipocytes were stained with 2 μg/mL Hoechst 33342 (ThermoFisher #H3570) and 2 μg/mL propidium iodide (ThermoFisher #P3566) in DMEM/F12 for 30 minutes at 37°C protected from light. To image floating adipocytes, an imaging chamber was created by stacking two 120 μm-thick imaging spacers (GraceBio #654004) on top of a microscope slide. Primary adipocytes were diluted in DMEM/F12, added as a 30 μL drop to the center of the spacers, and covered with a glass coverslip for imaging. The percentage of live adipocytes was quantified as the number of cells with propidium iodide-negative nuclei over the number of cells with Hoechst-positive nuclei. The imaging chamber was cleaned with 70% (v/v) ethanol, rinsed with 1x PBS, and dried between samples.

### RNA sequencing and analysis

For sample collection, 100 μL of packed adipocytes were lysed in 1 mL Qiazol (Qiagen #79306), incubated for 5 minutes at room temperature, and then stored at –80°C until RNA extraction. RNA extraction was performed using an RNeasy Lipid Tissue Mini Kit (Qiagen #74804) according to manufacturer protocols. RNA concentration, 260/280 purity, and 260/230 purity were assessed via a NanoDrop Spectrophotometer (ThermoFisher) and RNA concentration and quality were assessed via a 4200 TapeStation System (Agilent). Libraries were then prepared using a QuantSeq FWD Kit (Lexogen) and sequenced as 86 bp single-end reads on a NextSeq500 System (Illumina). Reads were trimmed using fastp (v.0.23.4) and aligned to the mouse reference genome GRCm39 (GENCODE) using STAR (v.2.7.10b). Reads were counted using the GeneCounts feature of STAR and differential expression analysis was performed with DESeq2 (v.1.40.2). Differentially expressed genes were defined as those with an absolute log_2_ fold-change greater than 1 and adjusted p-value less than 0.05. Heat maps were generated using pheatmap (v.1.0.12), volcano plots were generated using EnhancedVolcano (v.1.18.0), and principal component analysis plots were generated using PCAtools (v.2.12.0). Gene set enrichment analysis (GSEA) was performed using fgsea (v.1.26.0) by providing a list of ranked genes from the DESeq2 statistic column.

### Collection of mastectomy patient data

The Institutional Review Boards of Memorial Sloan Kettering Cancer Center (IRB 10-040) and Weill Cornell Medicine (IRB 1004010984-01) approved the collection of non-tumor breast tissue and blood under a biospecimen acquisition protocol. Informed consent was obtained from all participants. In total, 196 patients underwent mastectomy for breast cancer risk reduction or treatment. Height and weight were recorded prior to surgery and used to calculate body mass index (BMI). Standard definitions were used to categorize BMI as normal weight (BMI < 25), overweight (BMI 25.0–29.9), or obese (BMI ≥ 30). All data were reviewed for accuracy independently by research staff and a physician. On the day of mastectomy, paraffin blocks and snap frozen samples were prepared from breast WAT. If a tumor was present, samples were taken from an uninvolved quadrant of the breast. Two hematoxylin- and eosin-stained sections were generated from formalin-fixed, paraffin-embedded breast tissue in order to measure adipocyte diameters as previously described^40^. The sections were photographed with a 20x objective using an Olympus BX43 or BX50 microscope equipped with an Olympus DP27 or MicroFire (Optronics) digital camera, respectively. Mean diameters were calculated using measurements for each patient using the linear dimensional tool in the Canvas 11 Software (ACD Systems International, Inc.), which was calibrated using a microscope stage micrometer.

On the day of surgery, a 30 mL fasting blood sample was obtained preoperatively. Centrifugation was used to separate blood into serum and plasma within 3 hours of collection and stored at −80°C. Serum levels of triglycerides and high-density lipoprotein (HDL) cholesterol were determined in the Memorial Sloan Kettering Cancer Center Clinical Chemistry Lab.

Human RNA sequencing data have been previously published^10^ and were deposited to the European Genome-Phenome Archive (EGA) under accession number EGAS00001004665.

### Preparation of elastomer microwells

Elastomer microwells for *ex vivo* culture of primary adipocytes were prepared as previously described^9^. Briefly, SYLGARD 184 silicone elastomer (Ellsworth Adhesives #4019862) was cast in 1.5 mm-thick slabs according to manufacturer protocols. 10 mm outer diameter, 6 mm inner diameter elastomer rings were cut and bonded to glass coverslips using a plasma cleaner (Harrick Plasma). The rings were treated with 1% (v/v) poly(ethylenimine) (Sigma-Aldrich #181978), 0.1% (v/v) glutaraldehyde (ThermoFisher #ICN19859580), and washed three times with sterile water to facilitate covalent adhesion of adipocyte-collagen gels.

### Preparation of adipocyte-collagen gels

For *ex vivo* culture, packed adipocytes were mixed into a collagen gel at a 3:1 ratio (v/v) using wide-bore pipette tips as previously described^9^. Briefly, to prepare the collagen gel, collagen type I (VWR #47747-218) was diluted to a final concentration of 2.5 mg/mL in DMEM/F12. This solution was kept on ice and neutralized with 1N NaOH to begin polymerization immediately before the addition of packed adipocytes. During polymerization, the adipocyte-collagen gels were inverted every 2 minutes for 20 minutes at room temperature to ensure an even distribution of cells. The resulting gels were submerged in DMEM/F12 and cultured for up to 5 days at 37°C.

### Coculture studies

For coculture studies, 30,000 breast cancer cells (MDA-MB-231, MCF7, PY8119, or EO771) were plated per well in a 24-well plate containing 12 mm glass coverslips (Chemglass #CLS-1760-012) coated with 30 μg/mL collagen type I (VWR #47747-218). The following day, primary adipocytes were isolated and adipocyte-collagen gels in elastomer microwells were prepared. One cancer cell coverslip and one adipocyte-collagen gel were placed into each well of a 12-well plate along with 1.5 mL of DMEM/F12. Cancer cells and primary adipocytes were cocultured together for 72 hours. Cancer cells were then reserved for functional studies or fixed in 4% (w/v) paraformaldehyde for imaging.

### EdU proliferation assays

For proliferation studies, precultured cancer cells were trypsinized and reseeded at 30,000 cells per well in a 24-well plate containing 12 mm glass coverslips coated with 30 μg/mL collagen type I (VWR #47747-218). Once adherent, replated tumor cells were incubated in low-serum DMEM/F12 for 12 hours. The following morning, proliferating tumor cells were labeled with the Alexa Fluor 647 EdU Kit (ThermoFisher #C10340) according to manufacturer protocols. Briefly, a 20 μM EdU solution was prepared in low-serum DMEM/F12 and added 1:1 to existing media to achieve a final concentration of 10 μM EdU. Cancer cells were incubated for 2 hours at 37°C before proceeding with fixation, permeabilization, and detection of incorporated EdU. Cancer cells were co-stained with Hoechst 33342 (ThermoFisher #H3570) and BODIPY 493/503 (ThermoFisher #D3922). For inhibitor studies, 40 μM etomoxir (Millipore #236020) or DMSO vehicle were added with the low-serum DMEM/F12 after precultured cancer cells adhered to collagen-coated glass coverslips.

### Tissue-culture insert migration assays

For migration assays, precultured cancer cell coverslips were transferred to a 24-well plate and incubated in low-serum DMEM/F12 for 12 hours. The following morning, 8 μm pore tissue-culture inserts (VWR #29442-120) were coated with 30 μg/mL collagen type I (VWR #47747-218) and placed in 24-well plates containing 600 μL of DMEM/F12. Precultured cancer cells were then trypsinized and reseeded at 30,000 cells per tissue-culture insert in 100 μL of low-serum DMEM/F12 to establish a serum gradient. Cancer cells were allowed to migrate for 12 hours at 37°C. Cells were then fixed with 4% paraformaldehyde and a cotton swab was used to gently remove cells from the upper side of the insert membrane. The underside of each membrane was stained with DAPI (ThermoFisher #D1306) and imaged to quantify the number of transmigrated cancer cells. For inhibitor studies, 40 μM etomoxir (Millipore #236020) or DMSO vehicle were added to the low-serum DMEM/F12 when reseeding precultured cancer cells into tissue-culture inserts.

### Mitochondrial imaging in live breast cancer cells

To assess co-localization of breast cancer cell lipid droplets and mitochondria after coculture with adipocytes, precultured MDA-MB-231s were reseeded at 100,000 cells per glass-bottom petri dish (World Precision Instruments #FD35-100) coated with 30 μg/mL collagen type I (VWR #47747-218). Precultured cancer cells were then stained with 200 nM MitoView 640 (Biotium #70082) for 12 hours. Live samples were co-stained with 2 μg/mL Hoechst 33342 and 2.5 μg/mL BODIPY 493/503 for 30 minutes, washed twice with DMEM/F12, and imaged on an inverted Zeiss LSM 880 confocal microscope with an incubation chamber.

### Pre-labeling lipid to monitor transfer from adipocytes to breast cancer cells

For pre-labeling experiments, primary adipocytes were stained in suspension with 2.5 μg/mL BODIPY 493/503 for 30 minutes and washed three times with KRHB prior to embedding into collagen gels. Pre-labeled adipocytes were then cocultured with MDA-MB-231 breast cancer cells as described. After 48 hours, breast cancer cells were fixed with 4% (w/v) paraformaldehyde in 1x PBS and counterstained with 2.5 μg/mL DAPI and 165 μM Alexa Fluor 568 phalloidin. Breast cancer cells were then imaged on an inverted Zeiss LSM 880 confocal microscope to detect the presence of BODIPY-labeled neutral lipid.

### Sample fixation, permeabilization, and fluorescence staining

Samples were fixed in cold 4% (w/v) paraformaldehyde (Electron Microscopy Sciences #19208) in 1x PBS for 15 minutes at room temperature. Samples were washed twice with 1x PBS and permeabilized with 0.1% Triton X-100 (v/v) (ThermoFisher #A16046-AE) in 1x PBS supplemented with 1% (w/v) bovine serum albumin (BSA) (ThermoFisher #BP1600-100) for 15 minutes at room temperature. Samples were washed twice with 1x PBS and then stained with one or more of the indicated dyes as follows: 0.165 μM Alexa Fluor 488 phalloidin (ThermoFisher #A12379) or 0.165 μM Alexa Fluor 568 phalloidin (ThermoFisher #A12380) in 1x PBS with 1% BSA for 1 hour at room temperature; 2.5 μg/mL 4,4-difluoro-1,3,5,7,8-pentamethyl-4-bora-3a,4a-diaza-*s*-indacene (BODIPY 493/503) in 1x PBS with 1% BSA for 30 minutes at room temperature; 2.5 μg/mL 4’,6-diamidino-2-phenylindole, dihydrochloride (DAPI) (ThermoFisher #D1306) in 1x PBS with 1% BSA for 30 minutes at room temperature. Samples were then washed twice with 1x PBS and stored in 1x PBS at 4°C until imaged.

### Confocal microscopy and image analysis

Images were acquired on an inverted or upright Zeiss LSM 880 confocal microscope using either a 10x/0.45 W C-Apochromat objective, 20x/0.5 EC Plan-Neofluar objective, 20x/1.0 W Plan-Apochromat objective, 40x/1.2 W C-Apochromat objective, or 63x/1.4 O Plan-Apochromat objective. All images analysis was performed in QuPath^41^ and ImageJ (National Institutes of Health). For semi-automated quantification of adipocytes, cancer cell lipid droplets, and 3T3-L1 lipid droplets, the AdipoQ ImageJ plug-in^42^ was used. Manuscript figures were prepared using Adobe Photoshop, Adobe Illustrator, and Biorender.com.

### Substrate oxidation stress test

To assess ATP production and FAO, cancer cell metabolism was probed using a Seahorse XFe 96 Extracellular Flux Analyzer (Agilent) and Seahorse XF Long Chain Fatty Acid Oxidation Stress Test Kit (Agilent #103672-100). Control or precultured cancer cells were seeded at 20,000 cells per well in an XFe96 Cell Culture Microplate (Agilent #103794-100) in DMEM/F12 and allowed to adhere overnight. The following morning, media was changed to Seahorse XF DMEM, pH 7.4 (Agilent #103680-100), and manufacturer protocols were followed to complete the Long Chain Fatty Acid Oxidation Stress Test Kit. Inhibitor concentrations of 4 μM for etomoxir, 1.5 μM for oligomycin, 1 μM for FCCP, and 0.5 μM for rotenone/antimycin A were used. After the stress test was complete, DNA was extracted using lysis buffer (25 mM Tris-HCl, 0.4 M NaCl, 0.5% (w/v) sodium dodecyl sulfate) and total DNA content per well was measured using the QuantiFluor dsDNA Assay (VWR #PAE2671) for data normalization.

### Glycerol assay

Conditioned media from cancer cell and adipocyte-collagen gel cocultures was collected after 72 hours and stored at –20°C until processed. Once thawed, the relative degree of lipolysis during culture was estimated by measuring the concentration of glycerol in the conditioned media using the Adipolysis Assay Kit (Sigma-Aldrich #MAK313) according to manufacturer protocols.

### Isolation of extracellular vesicles from adipocyte-conditioned media

Adipocyte-collagen gels were cultured in 24-well plates containing 750 μL of exosome-depleted DMEM/F12 to generate conditioned media. Extracellular vesicles were then isolated from conditioned media using modified versions of published protocols^43^. Briefly, conditioned media was centrifugated at 300 x g at 4°C for 10 minutes to remove dead cells and other debris. The resulting supernatant was transferred to a new tube and centrifuged at 3000 x g at 4°C for 30 minutes to remove apoptotic bodies. The resulting supernatant was transferred to a new tube and combined at a 1:1 ratio (v/v) with 16% (w/v) PEG 6000 (Sigma-Aldrich #82160) in 1 M NaCl for a final concentration of 8% PEG 6000 in 0.5 M NaCl. The resulting solution was inverted several times and incubated at 4°C for at least 12 hours to precipitate soluble factors and extracellular vesicles. The precipitate was then pelleted via centrifugation at 3220 x g at 4°C for 1 hour. After aspirating the supernatant containing PEG 6000, the pellet was resuspended in 2 mL of filtered 1x PBS and ultracentrifuged in a TLS-55 swinging-bucket rotor (Beckman Coulter #346936) at 100,000 x g at 4°C for 4 hours. The supernatant containing soluble factors was discarded and the extracellular vesicle pellet was resuspended in 1 mL of filtered 1x PBS for nanoparticle tracking analysis.

### Nanoparticle tracking analysis of extracellular vesicles

Isolated extracellular vesicles were quantified via nanoparticle tracking analysis (NTA) using a Malvern NanoSight NS300. Samples were passed through the NanoSight flow cell using the included syringe pump at a flow rate of 50. At least 10 videos, each 30 seconds in length, were acquired with a 488 nm laser per sample. The concentration and size distribution of vesicles in each sample were then calculated using the Malvern Nanoparticle Tracking Analysis Software.

### Western blotting of vesicle markers

Sorted adipocytes for each condition, as well as preparations of the extracellular vesicles these cells generated, were lysed using lysis buffer (25 mM Tris, 100 mM NaCl, 1% Triton X-100, 1 mM EDTA, 1.0 μg/mL each of aprotinin and leupeptin, 1.0 mM β-glycerol phosphate, and 1.0 mM DTT). Protein concentrations of the cell (whole cell lysates; WCLs) and extracellular vesicle lysates were determined using the Bio-Rad Protein Assay Dye (Bio-Rad) and a spectrophotometer at a wavelength of 595. The lysates were normalized based on protein concentration and resolved on 4-20% SDS-PAGE gels (Invitrogen). The gels were transferred to PVDF membranes (Thermo-Fisher Scientific), and the membranes were blocked in 5.0% bovine serum albumin (BSA) in TBST buffer (20 mM Tris, 135 mM NaCl, and 0.02% Tween) for 1 hour at room temperature. The membranes were then incubated with one of the following primary antibodies diluted in TBST overnight at 4°C: FAK antibody (1:1000 dilution; Cell Signaling Technologies, #3285) and FLOT2 antibody (1:1000 dilution; Cell Signaling Technologies, #3436). The next day, the membranes were washed 3 times for 5 minutes with TBST before being incubated with HPR-conjugated secondary antibodies diluted in TBST for 1 hour at room temperature. The membranes were again washed with TBST 3 times for 5 minutes and exposed to ECL reagents (Bio-Rad). Images of the membranes were obtained using X-ray film (The Lab Depot).

### Breast cancer cell treatment with adipocyte-derived vesicles

For studies requiring treatment with adipocyte-derived extracellular vesicles, conditioned media was processed as described for analysis of vesicles through PEG 6000 precipitation and centrifugation at 3220 x g. The precipitate was then resuspended in 2 mL of exosome-depleted DMEM/F12 instead of 1x PBS. This solution was ultracentrifuged in a TLS-55 swinging-bucket rotor (Beckman Coulter) at 100,000 x g at 4°C for 4 hours. The resulting extracellular vesicle pellet was resuspended in 2 mL of exosome-depleted DMEM/F12 for downstream use.

### Extracellular vesicle treatment and DGAT1/2 inhibitor studies

For studies of lipid uptake from adipocyte-derived extracellular vesicles, 30,000 MDA-MB-231 breast cancer cells were plated per well in a 24-well plate containing 12 mm glass coverslips coated with 30 μg/mL collagen type I (VWR #47747-218). The following day, vesicles were applied to the cancer cells with 40 μM iDGAT1 (PF 04620110) (Tocris #1109276-89-2) and 40 μM iDGAT2 (PF 06424439) (Tocris #1469284-79-4) or an equivalent volume of DMSO vehicle. Treatment was applied such that each glass coverslip seeded with breast cancer cells received vesicles collected from one adipocyte-collagen gel over 24 hours. Breast cancer cells were incubated with vesicles for 24 hours at 37°C and then fixed with 4% (w/v) paraformaldehyde in 1x PBS for analysis of lipid uptake via fluorescence microscopy.

### Lipid extraction and derivatization for lipidomics

100 μL of flash-frozen adipocytes were lysed in 1 mL 66% methanol (v/v) in 0.1 M potassium phosphate buffer (pH 6.8) by vortexing twice on high for 20 seconds each. A 10 μL aliquot from each lysate was reserved for total protein quantification using BCA assay (Thermo Fisher, #23225). The remaining lysate was transferred to 13 x 100 mm borosilicate glass tubes (Fisher Scientific, #14-961-27), acidified with 10 μL of 1N HCl, and briefly vortexed. Total lipids were extracted twice with 1 mL of isooctane:ethyl acetate (3:1 v/v) and once with 1 mL hexanes. Each extraction was followed by 10-15 seconds of vortexing and then centrifugation at 2000 x g for 1 minute to ensure phase separation. The upper organic phases from all extractions were combined in fresh 13 x 100 mm borosilicate glass tubes, evaporated under gaseous nitrogen, and resuspended in 300 μL isooctane. For the non-esterified fatty acid (NEFA) fraction, 100 μL of the total lipid extract was transferred to a new glass tube, 25 ng internal stable isotope standard was added, evaporated under nitrogen, and resuspended in 25 μL of 1% pentafluorobenzyl bromide in acetonitrile (v/v). An equal volume of 1% diisopropylethylamine in acetonitrile (v/v) was added, followed by a 30-minute incubation at 25°C. The resulting pentafluorobenzyl-fatty acid derivatives were dried under nitrogen, resuspended in 100 µL of hexanes, and prepared for injection into the GC-MS. For the total fatty acid (TFA) fraction, 10 μL of the total lipid extract was transferred to a Teflon-lined glass tube, mixed with 75 ng of the stable isotope internal standard, and dried under nitrogen. Samples were resuspended in 500 μL of ethanol, and 500 μL of 1 M NaOH was added to saponify over 30 minutes at 90°C. After cooling, samples were acidified with 550 μL of 1 M HCl and extracted twice with 1.5 mL of hexanes. The extracts were dried under nitrogen and derivatized as described above. TFA samples were resuspended in 300 μL of hexanes for GC-MS injection.

### GC-MS conditions and data acquisition

For both the NEFA and TFA fractions, 1 μL of the pentafluorobenzyl-fatty acid derivatives was injected into an Agilent 8890 GC system coupled to a 5977B MS detector, using a DB-1MS UI column (Agilent, #122-0112UI). The GC temperature program consisted of an initial hold at 80°C for 3 minutes, followed by a ramp of 30°C per min to 125°C, and a second ramp of 25°C per min to 320°C with a final hold of 2 minutes. Methane was the carrier gas, flowing at 1.5 mL per min. Data were acquired in full-scan negative chemical ionization mode, using m/z for acyl chain lengths ranging from 8 to 22 carbons and unsaturation. Peak areas of analytes and internal standards were measured, and the ratio of analyte-derived ion areas to internal standard areas was calculated for quantification relative to each standard curve^44,45^.

### Statistical analysis

All experiments were performed with at least three biological replicates unless otherwise noted. Data with two conditions were evaluated with a nonparametric Mann-Whitney U test unless otherwise noted. Data with three or more conditions were evaluated with either a standard or nested one-way ANOVA with Tukey multiple comparisons test unless otherwise noted. P-values less than 0.05 were considered statistically significant. Unless otherwise noted, all data are plotted as means with standard deviations. All statistical analysis was performed using GraphPad Prism (v.10.2.3) or R (v.4.3.1).

