## Supplemental data for "Large adipocytes alter mode of lipid release and promote breast cancer malignancy"

### Supplemental figures

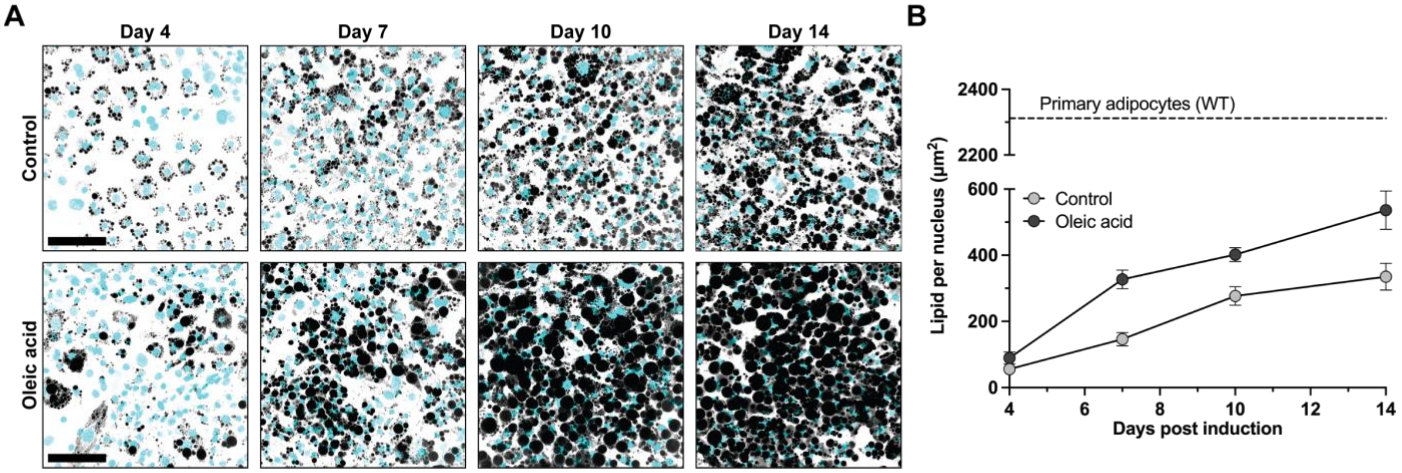


**Figure S1**. *In vitro* differentiation of adipocyte progenitor cells over time. (**A**) Representative images of 3T3-L1 preadipocytes at 4-, 7-, 10-, and 14-days following induction of adipogenic differentiation with or without oleic acid supplementation. Panels show cells stained for nuclei (cyan) and neutral lipid (gray). Scale bars = 100 µm. (**B**) Quantification of neutral lipid content per nucleus from (A) over time (n = 3). Dashed line represents the average lipid content of a wild-type primary adipocyte.

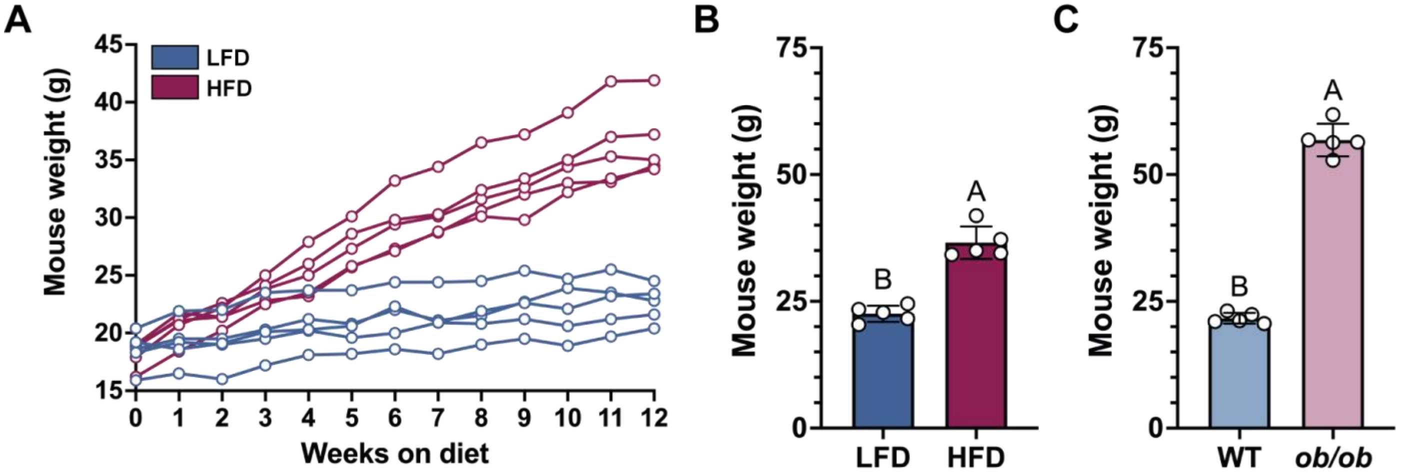


**Figure S2**. High-fat diet and leptin deficiency cause obesity in female C57BL/6 mice. (**A**) Mouse weights over 12 weeks for mice fed a low-fat diet (LFD) or high-fat diet (HFD) starting at 8 weeks of age. (**B**) Mouse weights at time of WAT collection for diet-induced obesity (n = 5). (**C**) Mouse weights at time of WAT collection for genetic obesity (n = 5). All data are presented as mean ± standard deviation unless otherwise noted. Statistics were performed with a Mann-Whitney U test. Compact letter display indicates statistical significance (p = 0.05).

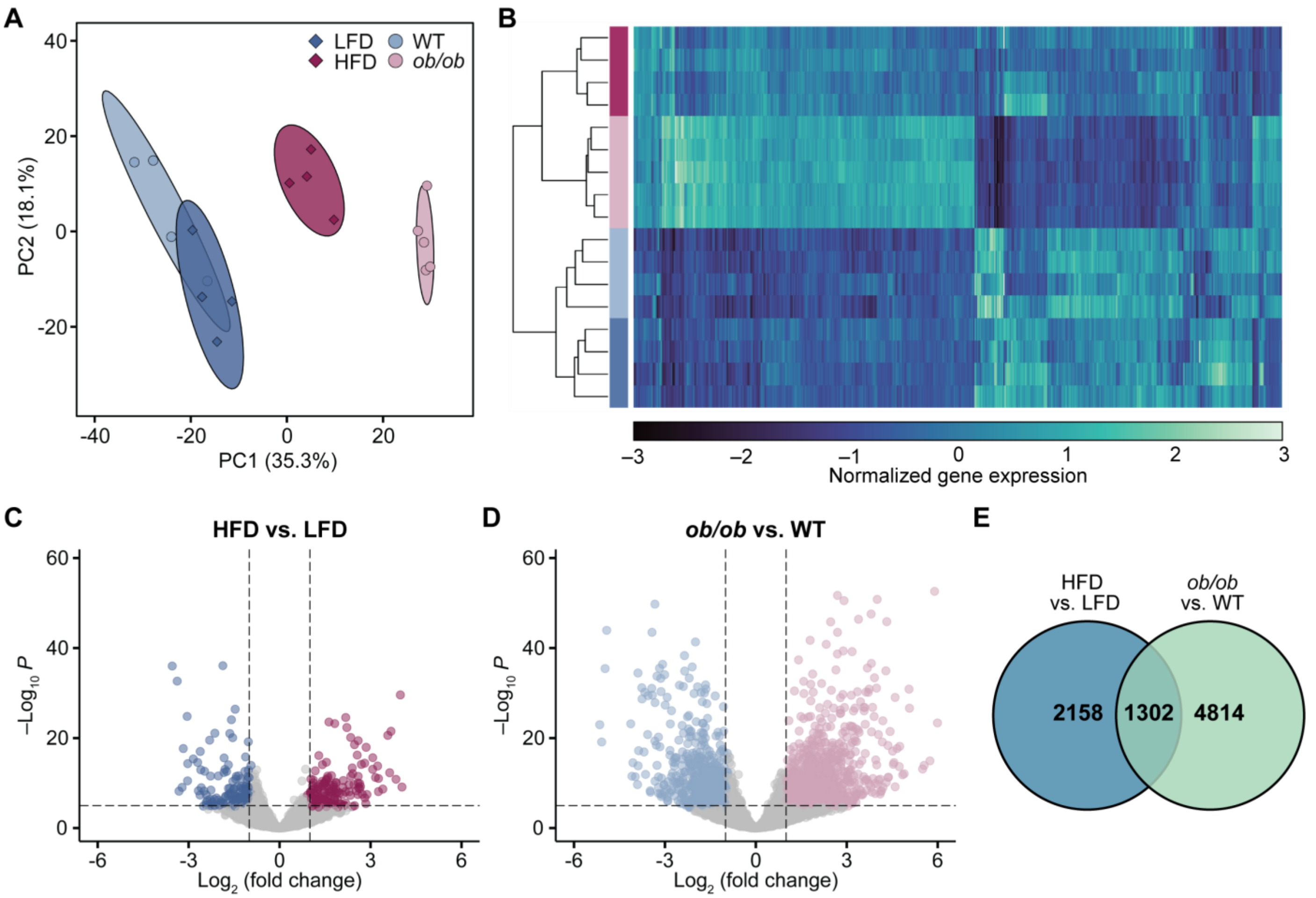


**Figure S3**. Transcriptional response of mammary adipocytes to diet-induced and genetic mouse models of obesity. (**A**) Principal component analysis of gene expression data for lean and obese adipocytes from both diet-induced and genetic mouse models of obesity (n = 4–5). Ellipses show the 95% confidence interval for each condition and individual points represent replicates (n = 4–5). (**B**) Heatmap with hierarchical clustering of normalized gene expression data for the top 500 most variable genes between samples. (**C**) Volcano plot of differentially expressed genes between adipocytes isolated from HFD and LFD mice. $Log_2$ (fold change) cutoff = 1.0, adjusted p-value cutoff = $1 \times 10^{-6}$. (**D**) Volcano plot of differentially expressed genes between adipocytes isolated from *ob/ob* and WT mice. $Log_2$ (fold change) cutoff = 1.0, adjusted p-value cutoff = $1 \times 10^{-6}$. (**E**) Venn diagram of differentially expressed genes (FDR < 0.05) between lean and obese adipocytes from diet-induced and genetic mouse models of obesity, including 1302 genes shared across both models.

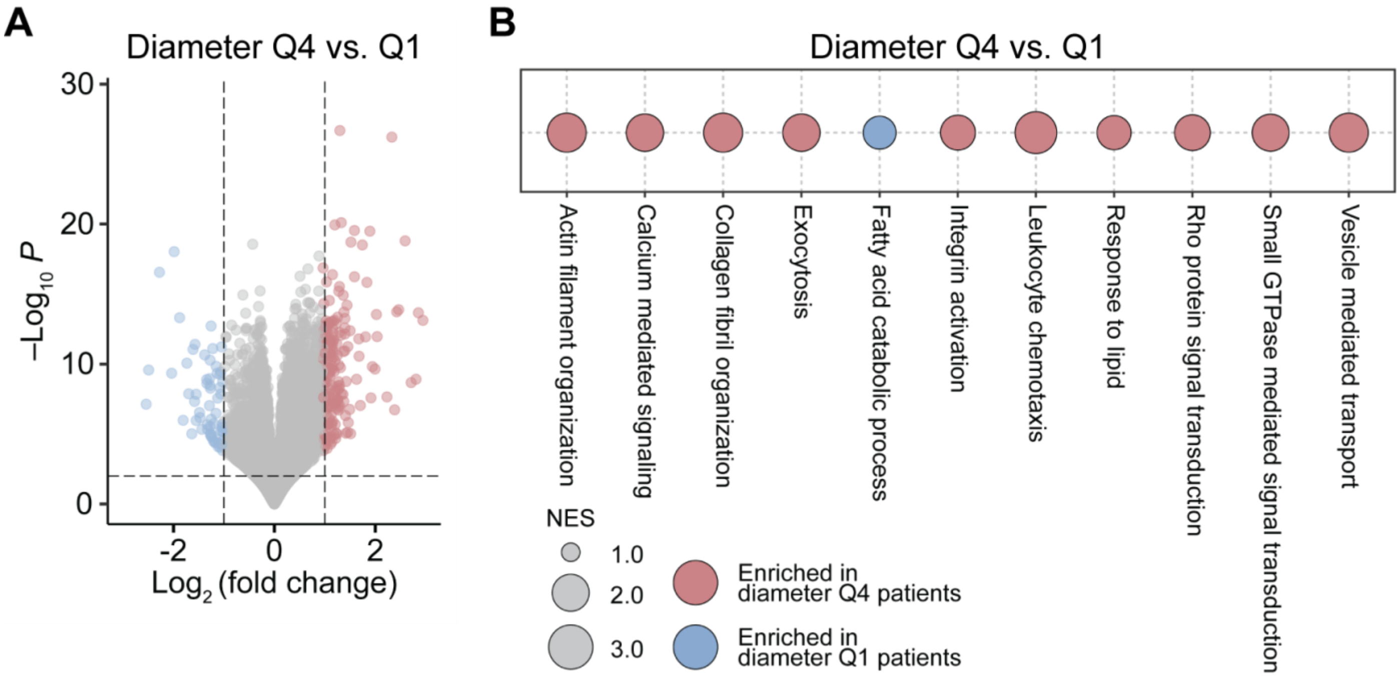


**Figure S4**. Differential expression and gene set enrichment analysis of human adipose tissue. (**A**) Volcano plot of differentially expressed genes between patients in Q4 and Q1. $Log_2$ (fold change) cutoff = 1.0, adjusted p-value cutoff = 0.05. (**B**) Gene sets from Gene Ontology Biological Processes (GOBP) enriched between patients in Q4 versus Q1. All comparisons shown meet the criteria set for statistical significance (FDR < 0.05).

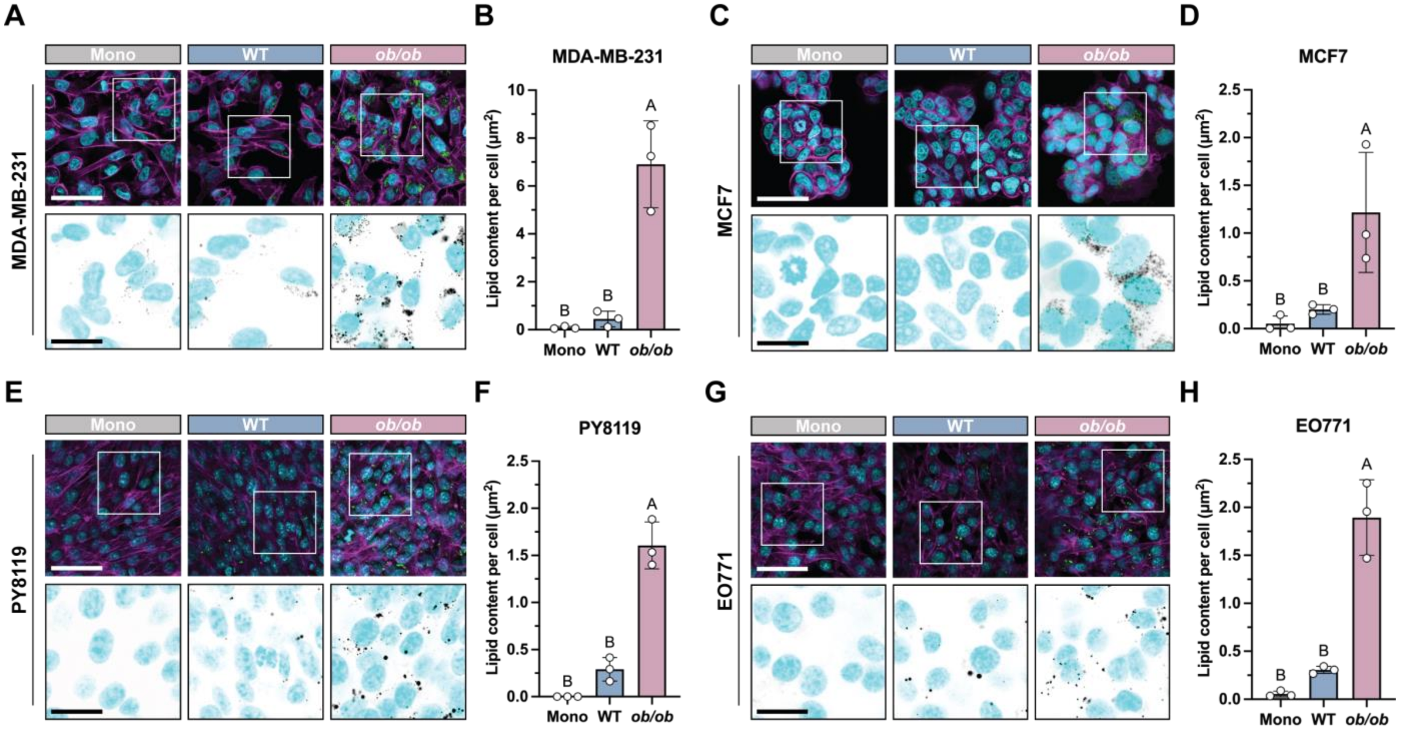


**Figure S5**. Adipocytes from genetically obese mice promote lipid accumulation in multiple mouse and human breast cancer cell lines. Representative images of MDA-MB-231 human breast cancer cells after monoculture (mono) or coculture with lean (WT) or unsorted obese (*ob/ob*) adipocytes for 48 hours. Scale bar = 100 µm. Upper panels show nuclei (cyan), neutral lipid (green), and actin filaments (magenta). Insets show nuclei (cyan) and neutral lipid (gray). Inset scale bar = 35 µm. (**B**) Quantification of neutral lipid staining per cancer cell (n = 3). Representative images and quantification are also provided for (**C–D**) MCF7 human breast cancer cells (n = 3), (**E–F**) PY8119 mouse breast cancer cells (n = 3), and (**G–H**) EO771 mouse breast cancer cells (n = 3). All data are presented as mean ± standard deviation unless otherwise noted. All statistics were performed with a one-way analysis of variance (ANOVA) with Tukey's multiple comparisons test. Compact letter display indicates statistical significance (p = 0.05).

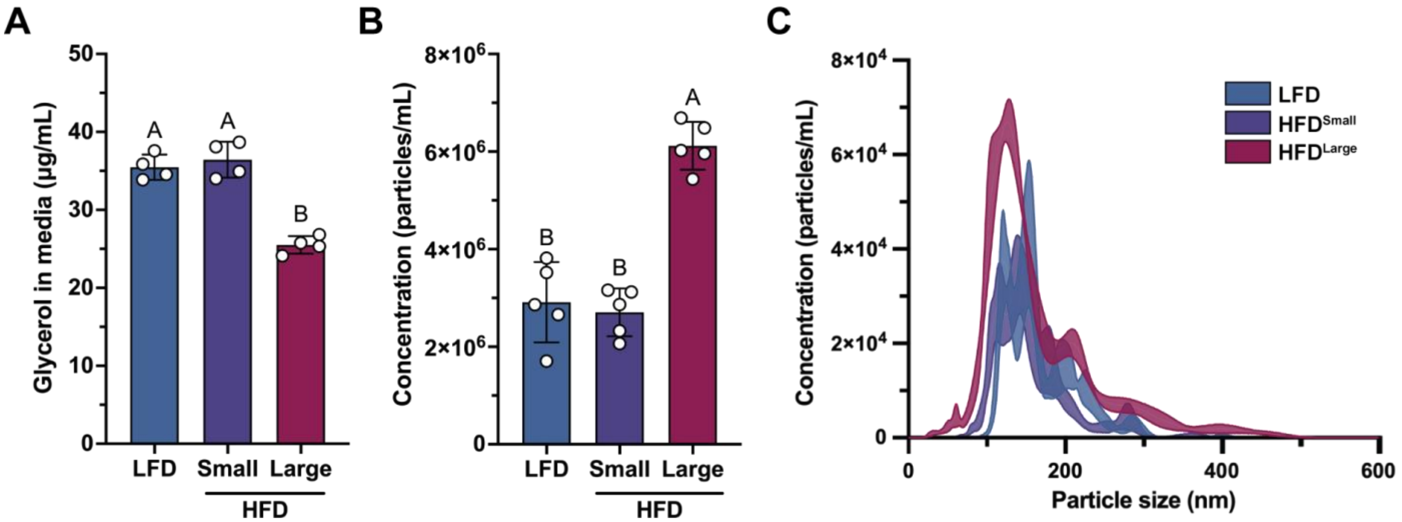


**Figure S6**. Large versus small adipocytes from high-fat diet mice secrete less glycerol and more extracellular vesicles. **A**) Concentration of glycerol in conditioned media collected from LFD or size-sorted HFD adipocytes in coculture with MDA-MB-231 breast cancer cells after 72 hours (n = 4). (**B**) Concentration of extracellular vesicles collected from LFD or size-sorted HFD adipocytes in monoculture after 24 hours (n = 5). (**C**) Size distribution of extracellular vesicles from (B) (n = 5). All data are presented as mean ± standard deviation. Statistics were performed with a one-way analysis of variance (ANOVA) with Tukey's multiple comparisons test. Compact letter display indicates statistical significance (p = 0.05).

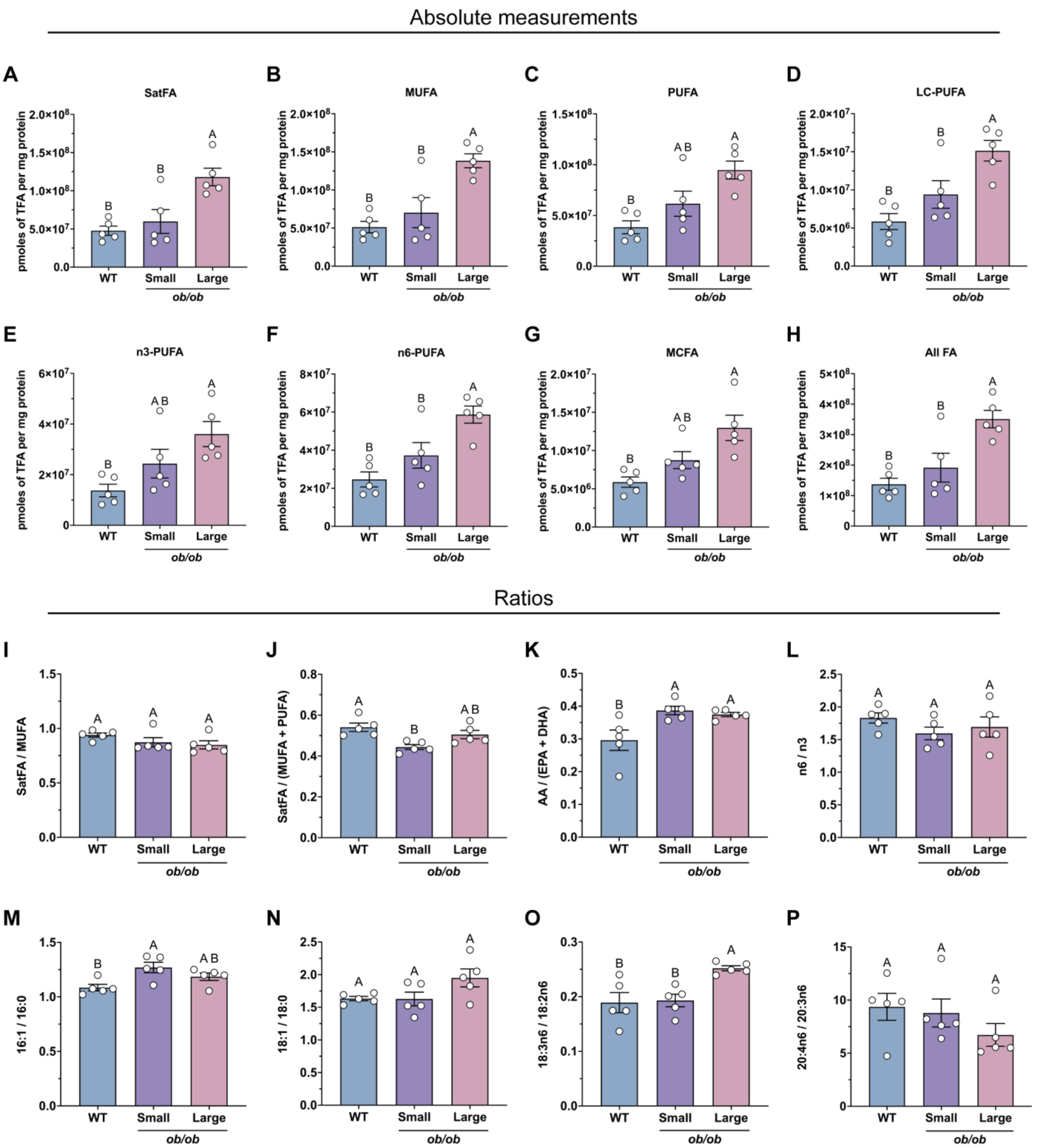


**Figure S7**. Lipidomics of total fatty acids from size-sorted obese adipocytes. Absolute abundance for total FAs normalized to mg protein for (**A**) saturated FAs, (**B**) monounsaturated FAs, (**C**) polyunsaturated FAs, (**D**) long-chain polyunsaturated FAs, (**E**) n3 polyunsaturated FAs, (**F**) n6 polyunsaturated FAs, (**G**) medium-chain FAs, and (**H**) all FAs (n = 5). Relative abundance for total FAs for (**I**) saturated versus monounsaturated FAs, (**J**) saturated versus monounsaturated and polyunsaturated FAs, (**K**) arachidonic versus eicosapentaenoic and docosahexaenoic acids, (**L**) n6 versus n3 polyunsaturated FAs, (**M**) palmitoleic versus palmitic acid, (**N**) oleic versus stearic acid, (**O**) gamma-linolenic versus linoleic acid, and (**P**) arachidonic versus dihomo-gamma-linolenic acid (n = 5). All data are presented as mean ± standard deviation. Statistics performed with a one-way analysis of variance (ANOVA) with Tukey's multiple comparisons test. Compact letter display indicates statistical significance (p = 0.05).

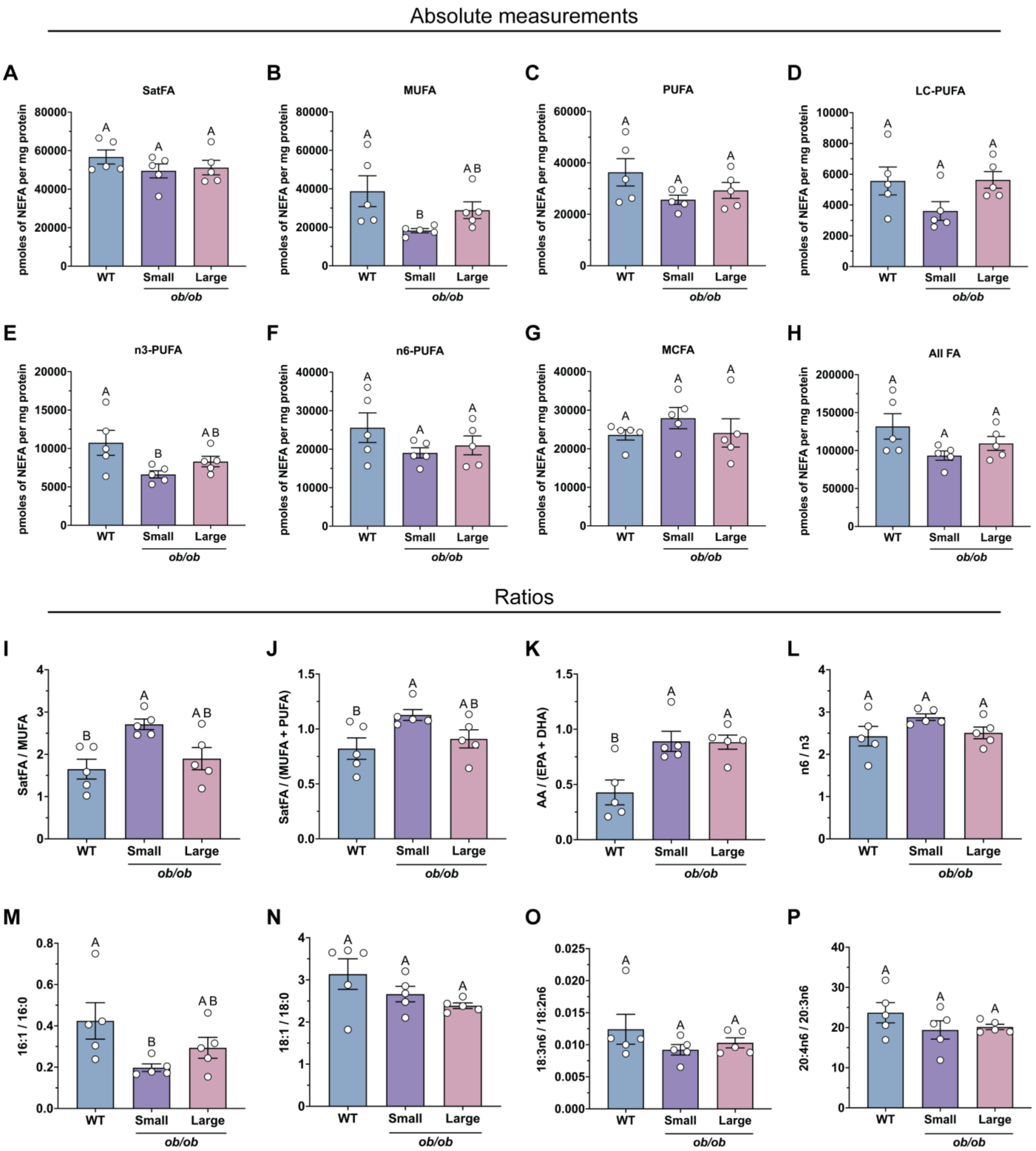


**Figure S8**. Lipidomics of non-esterified fatty acids from size-sorted obese adipocytes. Absolute abundance of non-esterified FAs normalized to mg protein for (**A**) saturated FAs, (**B**) monounsaturated FAs, (**C**) polyunsaturated FAs, (**D**) long-chain polyunsaturated FAs, (**E**) n3 polyunsaturated FAs, (**F**) n6 polyunsaturated FAs, (**G**) medium-chain FAs, and (**H**) all FAs (n = 5). Relative abundance of non-esterified FAs for (**I**) saturated versus monounsaturated FAs, (**J**) saturated versus monounsaturated and polyunsaturated FAs, (**K**) arachidonic versus eicosapentaenoic and docosahexaenoic acids, (**L**) n6 versus n3 polyunsaturated FAs, (**M**) palmitoleic versus palmitic acid, (**N**) oleic versus stearic acid, (**O**) gamma-linolenic versus linoleic acid, and (**P**) arachidonic versus dihomo-gamma-linolenic acid (n = 5). All data are presented as mean ± standard deviation. Statistics performed with a one-way analysis of variance (ANOVA) with Tukey's multiple comparisons test. Compact letter display indicates statistical significance (p = 0.05).
